# Genomic insights into longan evolution from a chromosome-level genome assembly and population genomics of longan accessions

**DOI:** 10.1101/2021.05.27.445741

**Authors:** Jing Wang, Jianguang Li, Zaiyuan Li, Bo Liu, Lili Zhang, Dongliang Guo, Shilian Huang, Wanqiang Qian, Li Guo

## Abstract

Longan (*Dimocarpus longan*) is a subtropical fruit best known for its nutritious fruit and regarded as a precious tonic and traditional medicine since ancient times. High-quality chromosome-scale genome assembly is valuable for functional genomic study and genetic improvement of longan. Here, we report a chromosome-level reference genome sequence for longan cultivar JDB with an assembled genome of 455.5 Mb in size anchored to fifteen chromosomes, representing a significant improvement of contiguity (contig N50=12.1 Mb, scaffold N50= 29.5 Mb) over a previous draft assembly. A total of 40,420 protein-coding genes were predicted in *D. longan* genome. Synteny analysis suggests longan shares the widespread gamma event with core eudicots, but has no other whole genome duplications. Comparative genomics showed that *D. longan* genome experienced significant expansions of gene families related to phenylpropanoid biosynthesis and UDP-glucosyltransferase. Deep genome sequencing analysis of longan cultivars identified longan biogeography as a major contributing factor for genetic diversity, and revealed a clear population admixture and introgression among cultivars of different geographic origins, postulating a likely migration trajectory of longan overall confirmed by existing historical records. Finally, genome-wide association studies (GWAS) of longan cultivars identified quantitative trait loci (QTL) for six different fruit quality traits and revealed a shared QTL containing three genes for total soluble solid and seed weight. The chromosome-level reference genome assembly, annotation and population genetic resource for *D. longan* will facilitate the molecular studies and breeding of desirable longan cultivars in the future.

## INTRODUCTION

Longan (*Dimocarpus longan* Lour.), also known as dragon’s eyeball and closely related to lychee, is a tropical/subtropical evergreen fruit tree in Sapindaceae family with a diploid genome^1^ (2n = 2x = 30). It is regarded as a precious tonic and traditionally used as a medicinal plant with rich pharmaceutical effects from many parts of the plant, mainly fruits, contributing to the rural economic development in tropical and subtropical areas. The usage of longan in traditional herbal remedies was recorded in the compendium of materia medica (Ben Cao Gang Mu in Chinese) by Li Shizhen, a famous traditional Chinese medicine expert during the Ming Dynasty, who called longan the king of fruits ^2^. Given its high nutritional and economic value, longan is cultivated in many countries such as China, Australia, Thailand and Vietnam. Among these countries, China has the highest production of longan with largest cultivation area^3^ including Guangdong, Guangxi, Fujian, Hainan and other regions in China^4^.

China has a long history of longan cultivation and uses. Native to South China, longan has been cultivated in China for over 2000 years with rich germplasm resources^5^. Abundant wild resources are found in Yunnan and Hainan province, where longan was introduced to South Asian countries such as Thailand^6^. A previous study based on the different pollen exine patterns of fourteen longan varieties supports Yunnan as the primary center of longan origin, and Guangdong, Guangxi and Hainan as the secondary centers^7^. To date the population genetic structure of longan remains elusive. Longan varieties have rather ambiguous genetic background due to the fact that they reproduce via both inbreeding and crossbreeding in the field. A previous analysis using ISSR (Inter-simple sequence repeat) markers indicated that Thailand and Vietnam varieties have close genetic relationships^8^. However, the classification of longan varieties using these markers has been inconclusive due to marker selection, number of varieties and classification methods being adopted. A resolved population structure of longan varieties and better understanding of its genetic diversity and migration history is essential for conservation and breeding of longan but remains limited, requiring a large-scale phylogenomic study of longan varieties around China and Southeast Asia.

Longan leaf, flower, fruits, and seeds^9, 10^ are rich in polyphenols with anti-cancer, anti-oxidant properties, biosynthesized primarily through the shikimic acid, flavonoid and phenylpropanoid pathways. Phenylpropanoid pathway is one of the most extensively investigated specialized metabolic pathways. The branches of phenylpropanoid pathway produce metabolites such as flavonoids, hydroxy-cinnamic acid esters, hydroxycinnamic acid amides, and the precursors of lignin, lignans, and tannins. These phenolic natural products are key components of nutritious, flavor and medicinal properties in longan. Phenylpropanoids also play important roles in plant resistance to pathogen infection^11^ either by acting as physical barriers against invasion, or through chemical toxicity to herbivores and microbial pathogens. Therefore, these secondary metabolites are acquired traits that on one hand offer adaptative advantage to longan through evolution, and on the other hand can be exploited by humankind as useful medicines. Despite the vital roles of phenylpropanoids to the nutrition and flavor of longan fruits, the biosynthetic pathways for these metabolites in longan remain uncharacterized due to limited genetic and genomic resources and technical difficulty of genetic transformation, impeding the improvement of desired traits in longan fruits through molecular and genomic breeding.

Longan breeding, mainly relying on sexual hybridization, typically targets two main traits: fruit size and sweetness^12^. It is challenging and time-consuming to do longan breeding due to its long juvenile period and difficulty of genetic transformation. Marker-assisted selection (MAS) is an effective biotechnological tool to promote early selection of hybrid progenies at seedling stage^13^. Yet our knowledge about genetic mapping of longan is limited. Guo *et al*. (2011) constructed a low-quality male and female genetic map, consisting of 243 and 184 molecular markers separately^14^. Single nucleotide polymorphism (SNP) markers based on restriction site associated DNA sequencing (RAD-seq) was constructed for QTL identification by using hybrid progenies F_1_ and two parents as materials^13^. A high-quality reference genome sequence and knowledge of the longan genetic background would significantly facilitate the investigation of genotype-phenotype association of longan germplasms and thus expedite the longan breeding program. Although a draft genome sequence of *D. longan* “HHZ” cultivar is available^9^, the assembly is essentially fragmented composed of 51,392 contigs with a contig N50 of 26kb.

Here, we produced a chromosome-level genome assembly for *D. longan* JDB cultivar combining Illumina paired-end (PE), PacBio single molecule real-time (SMRT) sequencing and high-throughput chromatin conformation capture sequencing (Hi-C). We annotated the genome using *ab inito* prediction, homolog evidence and multi-tissue transcriptomic data. In addition, we conducted population genome sequencing from a collection of longan accessions, followed by an in-depth analysis of population structure using high-quality genetic variants. The analysis revealed the population genetic diversity of longan and demonstrated the population admixture and introgression among cultivars from major longan growing areas. GWAS analysis revealed three genes were associated with fruit total soluble solid and seed weight, suggesting parallel evolution in the evolutionary diversification of domesticated species. The genome assembly, annotations and genetic variants was valuable to functional genomic studies as well as molecular breeding of *D. longan* for improving the yield, fruit quality and exploiting its medicinal properties.

## RESULTS

### Genome assembly and annotation

*D. longan* “JDB” cultivar originated from Fujian is planted in Longan Germplasm Repository of Guangdong Province (Figure 1A-1D), and the fresh young leaves were collected for genomic DNA isolation and sequencing. To construct a chromosome-level reference genome for *D. longan*, a total of 184.4Gb PacBio SMRT reads (∼415x coverage), 25.3Gb (∼56x coverage) Illumina PE reads, and 57.6Gb (∼127x coverage) Hi-C Illumina read pairs were generated in this study (Supplementary Table 1). We estimated the genome size of *D. longan* cultivar JDB as 474.98 Mb with a heterozygosity rate of 0.36% via the distribution of k-mer frequency using Illumina PE reads (Figure 1E). PacBio SMRT reads were used to assemble the *D. longan* genome by *Canu*^15^, followed by polishing contigs using Illumina PE reads by *Pilon*^16^, which yielded a draft genome assembly of 455.5Mb (Table 1). Next, Hi-C paired-end reads were used to anchor the contigs to chromosomes with *3D-DNA*^17^. The final *D. longan* JDB genome assembly of 455.5Mb covers 95.90% of the estimated genome size (474.98Mb) and 98.7% of sequences were anchored onto 15 chromosomes (Figure 1F) with contig and scaffold N50 of 12.1Mb and 29.6Mb, respectively (Table 1). Thus, this longan genome assembly represents a significant improvement over the highly fragmented *D. longan* HHZ genome assembly (contig N50: 0.026 Mb) previously released^9^. Genome completeness was assessed using the plant dataset of the Benchmarking Universal Single Copy Orthologs (BUSCO) database v1.22^18^, with e-value < 1e−5. BUSCO evaluation revealed the completeness of 98.1% for our *D. longan* genome assembly (88.4% single copy; duplicated copy 9.7%, 1.1% fragmented and 0.8% missing) (Table 1, Supplementary Table 2).

**Figure 1:**
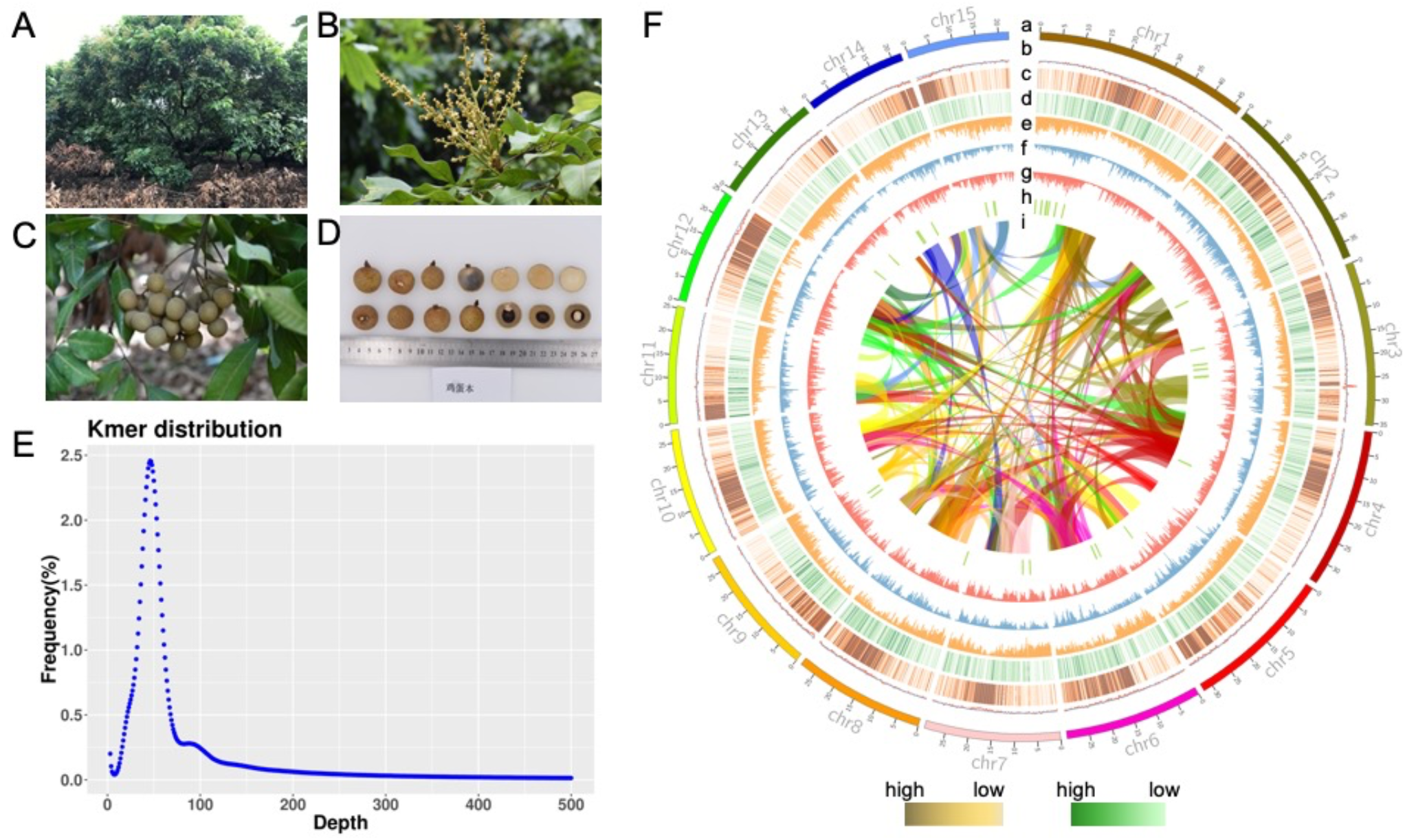
Chromosome-level genomic assembly of longan (*Dimocarpus longan* Lour.). (**A-D**): Photos of flower (**B**), fruit cluster (**C**), and fruit section (**D**) of longan cultivar JDB. (**E**) Kmer frequency distribution analysis for JDB genome based on Illumina paired-end reads. (**F**) Overview of *D. longan* genome. Track a to i: chromosomes, GC-content, density of *Gypsy* LTR (long terminal retrotransposons), density of *Copia* LTR, density of protein-coding genes, SNP density, Indel density, distribution of secondary metabolic gene cluster (predicted using *plantismash*), syntenic blocks (color ribbons). The density statistics is calculated within genomic windows of 150kb in size.

**Table 1.** Statistics for *Dimocarpus longan* JDB genome assembly and annotations.

|  | Statistics | *D. longan* JDB<br>(this study) | *D. longan* Honghezi<sup>9</sup> |
| --- | --- | --- | --- |
| **Contig** | Total number of contigs | 250 | 51,392 |
|  | Assembly size (Mb) | 455.5 | 471.9 |
|  | Contig N50 (Mb) | 12.1 | 0.026 |
|  | Contig N90 (Mb) | 1.8 | 0.006 |
|  | Largest Contig (Mb) | 31.1 | 0.17 |
| **Scaffold** | Total number of scaffolds | 90 | 17,367 |
|  | Assembly size (Mb) | 455.5 | 495.3 |
|  | Scaffold N50 (Mb) | 29.6 | 0.57 |
|  | Scaffold N90 (Mb) | 22.3 | 0.12 |
|  | Largest scaffold (Mb) | 46.6 | 6.9 |
| **Annotation** | Number of genes | 40,420 | 31,007 |
|  | Repeat content (%) | 41.7 | 52.9 |
|  | Number of ncRNA | 2,555 | NA |
|  | BUSCO (%) | 98.1% | 94% |
|  | GC content (%) | 43.9 | 33.7 |

We next performed genome annotations by using the *BRAKER2* pipeline combining evidence from *ab initio* prediction, protein homologs and multi-tissue (root, shoot, leaf and fruit) transcriptome sequencing data. The genome annotation pipeline predicted a total of 40,420 protein-coding genes and 2,555 non-coding RNAs for *D. longan*, respectively (Table 1). Longan genome has an overall guanine-cytosine (GC) content of 34 % and gene density of 89 genes per Mb (Supplementary Table 2). About 89.0 % genes have been annotated with NR (non-redundant protein sequence database) and 84.6 % genes with KEGG (Kyoto encyclopedia of genes and genomes) terms (Supplementary Table 3). Repetitive elements make up 41.7 % of *D. longan* genome, of which 54.9% and 25.4 % are long terminal repeat retrotransposons (LTRs) and DNA transposons respectively. Two major LTR subtypes, LTR-*Copia* (179.64 Mb) and LTR-*Gypsy* (66.18 Mb) represent 8.55 % and 15.53 % of the longan genome, respectively (Supplementary Table 4).

### Comparative genomics and synteny analysis

Next, we performed intraspecies synteny analysis of *D. longan* genome to investigate its genome evolution history. Intraspecies syntenic gene pairs in *D. longan* were identified using *MCScanX*, which supported the presence of a whole genome triplication (WGT) event in longan genome (Figure 2A). Distribution of synonymous substitution rate (*Ks*) for the syntenic gene pairs also supported that the *D. longan* genome experienced the WGT (Figure 2B). In addition, the l:1 ratio of syntenic blocks between longan and grape (*Vitis vinifera*) and 1:2 ratio of syntenic blocks between longan and poplar (*Populus trichocarpa*) indicated the longan only had WGT (γ) event, and did not have other whole genome duplications (WGDs) (Figure 2C). To reveal the genome evolution and divergence of longan, we performed phylogenomic analysis of longan and thirteen representative plant species including eight Rosids (*Citrus sinensis, Carica papaya, Arabidopsis thaliana, Theobroma cacao, P. trichocarpa, Ricinus communis, Glycine max, V. vinifera*), two Solanaceae *(Solanum tuberosum, Nicotiana attenuata)*, one Poaceae *(Oryza sativa)* and a basal angiosperm *(Amborella trichopoda)*. Orthogroup (gene family) identification revealed that these plants shared 7530 orthogroups, 137 of which are single-copy ones (Figure 3A; Supplementary Table 5). Particularly, we identified 1366 orthogroups unique to *D. longan* comparing to *A. thaliana, C. cinensis, S. tuberosum* and *P. trichocarpa* (Figure 3B). The multiple sequence alignment of 137 single-copy orthologs in 14 species were concatenated and used for phylogeny construction followed by a divergence time estimation using *MCMCTREE* calibrated with fossil record time (Figure 3C). We found that among the thirteen species, longan was phylogenetically closest to *C. sinensis*. The two species shared a last common ancestor at around 67 million years ago (Mya) that diverged from asteroids (*N. attenuata, S. tuberosum*) at around 125 Mya (Figure 3C).

**Figure 2.**
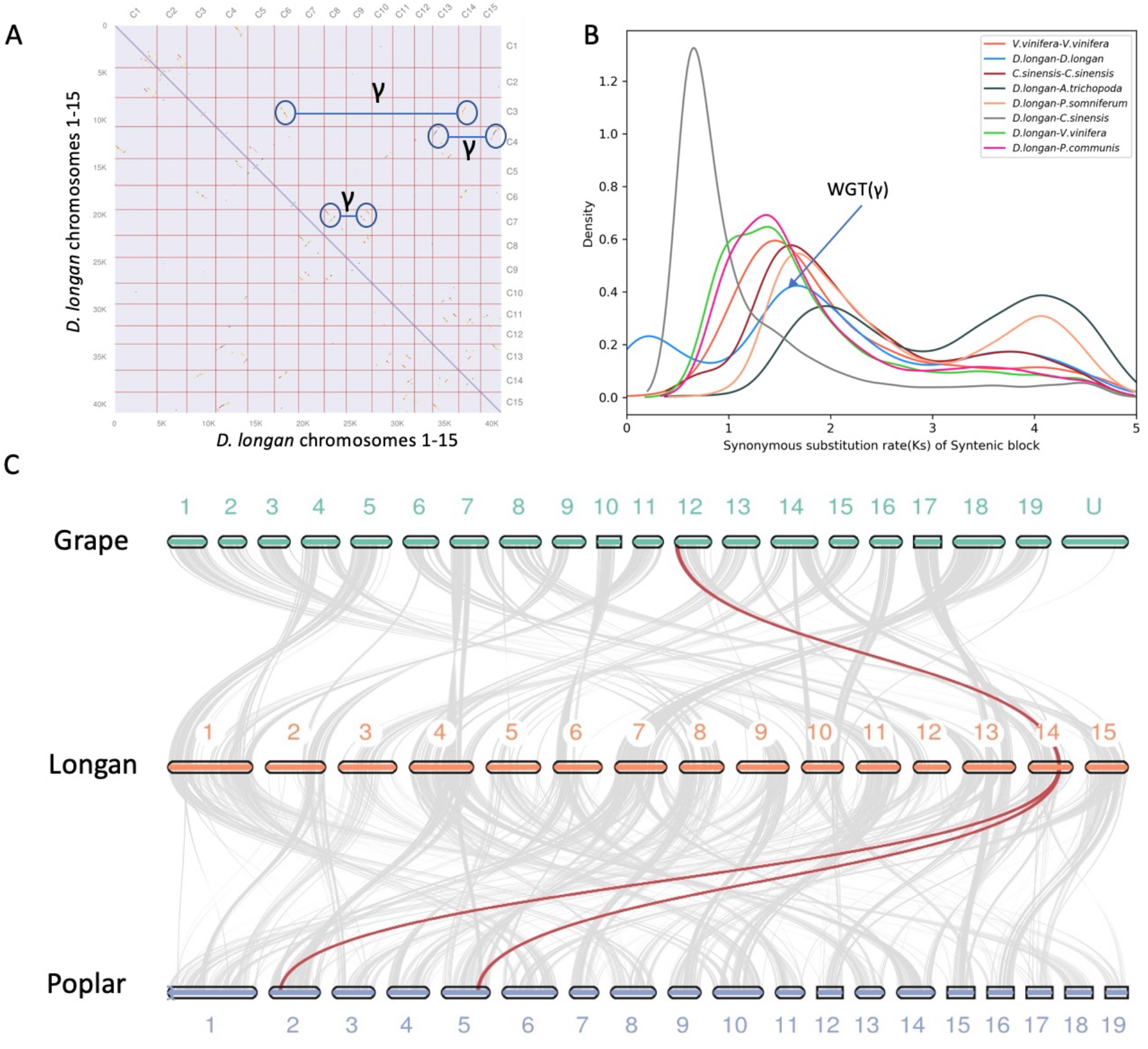
Comparative genomics and synteny analysis of *Dimocarpus longan*. (**A)** Whole genome dot plot of *D. longan* showing intraspecies genome synteny based on syntenic gene pairs. The pair of black circles connected by a straight line highlight the syntenic blocks detected in *D. longan* genome, which corresponds to the whole genome triplication (γ event). (**B**) Distribution of *Ks* (synonymous substitution rate) density for syntenic paralogs or orthologs detected in pairwise comparisons among various plant genomes. **(C)** Karyotype macrosynteny plots displaying the collinear relationships for different chromosomes among grape (*Vitis vinifera*), longan (*Dimocarpus longan*) and poplar (*Populus trichocarpa*). The colored lines highlight the syntenic blocks conserved among three species.

**Figure 3.**
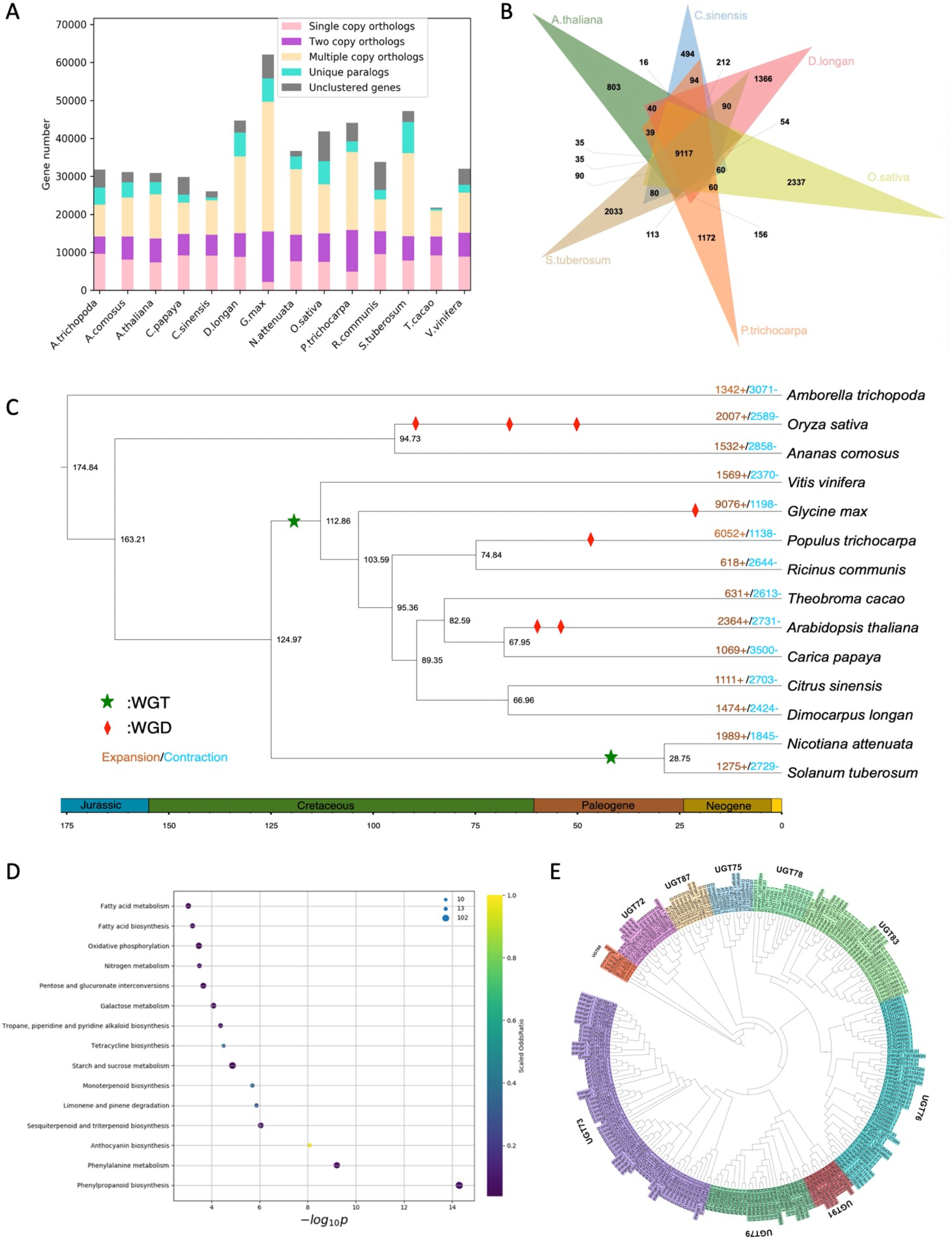
Phylogenomic genomics of *Dimocarpus longan*. (**A)**. Summary of gene family clustering of *D. longan* and 13 related species. Single copy orthologs: 1-copy genes in ortholog group. Multiple copy orthologs: multiple genes in ortholog group. Unique orthologs: species-specific genes. Other orthologs: the rest of the clustered genes. Unclustered genes: number of genes out of cluster. (**B)**. Comparison of orthogroups (gene families) among six angiosperm species including *D. longan* (longan), *A. thaliana* (Arabidopsis), *C. sinensis* (citrus), *S. tuberosum* (potato), *P. trichocarpa* (poplar) and *O. sativa* (rice). (**C)**. Phylogenetic relationship and divergence time estimation. The number of gene family expansion and contraction was indicated by red and blue number, respectively. (**D)**. Bubble plot summarizing the most significantly enriched KEGG (Kyoto Encyclopedia of Genes and Genomes) terms associated with *D. longan* expanded gene families. X-axis is the log10 transformed p-value. The size of bubble is scaled to the number of genes. The color scale represents the scale of odds ratio in observed versus expected (genomic background) number of genes annotated with specific KEGG terms. (**E)**. A phylogenetic tree of UGTs (UDP-glucosyltransferase) in three angiosperms including *D. longan*.

### Expanded gene families related to phenylpropanoid biosynthesis and UDP-glucosyltransferases

Gene family contraction and expansion are the evolutionary forces that drive the rapid speciation and result in the diversification of plants^9^. Gene family analysis suggested longan genome has 1474 expanded and 2424 contracted gene families (Figure 3C) compared with common ancestor of *C. sinensis* and *D. longan*. KEGG (Kyoto Encyclopedia of Gene and Genomes) enrichment showed that expanded gene families were significantly enriched (*P < 0*.*05*) with “phenylpropanoid biosynthesis”, “phenylalanine metabolism”, “anthocyanin”, “sesquiterpenoid and triterpenoid biosynthesis”, “monoterpenoid biosynthesis” (Figure 3D) The 97 expanded longan phenylpropanoid biosynthesis genes were classified into seven gene families: phenylalanine ammonia-lyase (PAL, 5 members), peroxidase (POD, 38 members), O-methyltransferase (OMT, 3 members), glycosyl hydrolase family 1 (GH1, 26 members), aldehyde dehydrogenase family (ADH, 18 members) and AMP-binding enzyme (4 members), beta-galactosidase (BGL, 3 members) (Supplementary Table 6), likely involved in the biosynthesis of various precursors to lignin. Lignin is a major component of certain plant cell walls and likely involved in the longan speciation^19^. Structural lignins provide physical barriers against pathogen infection, and mechanical support for plant growth and the long-distance transportation of water and nutrients^20^. Key enzymes such as PAL, POD and PPO of phenylpropanoid and lignin pathway were involved in these processes. Catalyzing the first step in the phenylpropanoid biosynthetic pathway, PALs were expressed at the higher levels in the roots, leaves and stems but not in green fruits (Supplementary Figure 1), consistent with a previous report^9^. In addition, 28 of the 38 PODs showed differential expression in four major tissues (leaves, stems, roots and fruits) (Supplementary Figure 2). A previous transcriptome-based study contradicted with our finding, suggesting that structural genes in phenylpropanoid pathways showed contraction instead of expansion^9^. However, we argued that a high-quality genome assembly and annotation is essential to a correct and complete characterization of gene families, therefore more trustworthy than transcriptome analysis alone.

Plants have evolved exquisite mechanisms for the biosynthesis of phenylpropanoids through acylation, methylation, glycosylation and hydroxylation^21^. Most of the compounds synthesized by the phenylpropanoid pathway can be glycosylated by UDP-glucosyltransferase (UGT). UGTs are key glycosylation enzymes that stabilize and enhance the solubility of small metabolites to maintain intracellular homeostasis^22^. For example, UGTs glycosylate volatile benzenoids/phenylpropanoids, monoterpene linalool, and a strawberry aroma 4-hydroxy-2,5-di-methyl-3(2H)-furanone etc. InterPro (IPR) protein domain enrichment analysis showed that the longan expanded gene families are significantly enriched with IPR domains such as UGTs and cytochrome P450s (Supplementary Figure 3). Longan genome encodes more (215) UGTs (Supplementary Table 7) than *Arabidopsis* (107), *C. grandis* (145) and *V. vinifera* (181) do, but fewer than in apple does (241) ^23^. UGTs also participate in plant development, growth and defense responses. Phenylpropanoid metabolism plays important roles in resistance to pathogen infection through the UGTs^21^. A phylogenetic tree was constructed using UGT protein sequences in longan and other plants including *Arabidopsis, Citrus* (Figure 3e), in which 115 expanded UGTs were divided into ten groups. Five groups A, D, E, G, and L expanded more than others, although the number of genes in these groups varied widely among species^23^. In longan, group A, D, H, I, and L expanded more than the other groups. For example, the number of longan UGTs in group D (31 UGTs, UGT73) and group I (19 UGTs, UGT83) was significantly increased compared to those in *Arabidopsis* and *Citrus*. A group D member UGT73C7 mediated the redirection of phenylpropanoid metabolism to hydroxycinnamic acids (HCAs) and coumarin biosynthesis under biotic stress, resulting in SNC1-dependent Arabidopsis immunity^24^. In group I, number of UGTs was highest compared to other fruits such as peach (5 UGTs), apple (11 UGTs) and grapevine (14 UGTs). UGT83A1 (GSA1) was required for metabolite reprogramming under abiotic stress through the redirection of metabolic flux from lignin biosynthesis to flavonoid biosynthesis and the accumulation of flavonoid glycosides, which coordinately confer high crop productivity and enhanced abiotic stress tolerance^25^. Transcriptomic profiling suggested 96 UGTs were differentially expressed in longan, with four (accounting for 4.2%, fourteen (14.6%), and ten (10.4%) UGTs were uniquely expressed in leaf, root and fruit, respectively (Supplementary Figure 4, Supplementary Table 8).

### Population structure, migration and genetic admixture of longan cultivars

To understand the longan genomic dynamics across its current distribution range in southern China and southeast Asian countries, we performed whole genome resequencing analysis of 87 accessions (Supplementary Table 9) from five southern provinces in China: Guangdong, Fujian, Guangxi, Sichuan, Hainan, and three other countries Thailand, Vietnam and Australia, with an average sequencing depth of 50×. Read mapping to longan reference genome and variant detection yielded 29,730,132 single nucleotide polymorphisms (SNPs). After filtering, 11,421,213 high-quality SNP loci (minor allele frequency > 5%) were used for subsequent population genetic analyses. Although Guangdong borders on Fujian (Figure 4A), the climate of the two provinces was largely different during longan growing season. After generations of planting and screening, different cultivation areas have formed their own longan variety characteristics and types. Using the genetic variant data, we analyzed the population structure within these longan cultivars using phylogenomic analysis and principal component analysis.

**Figure 4.**
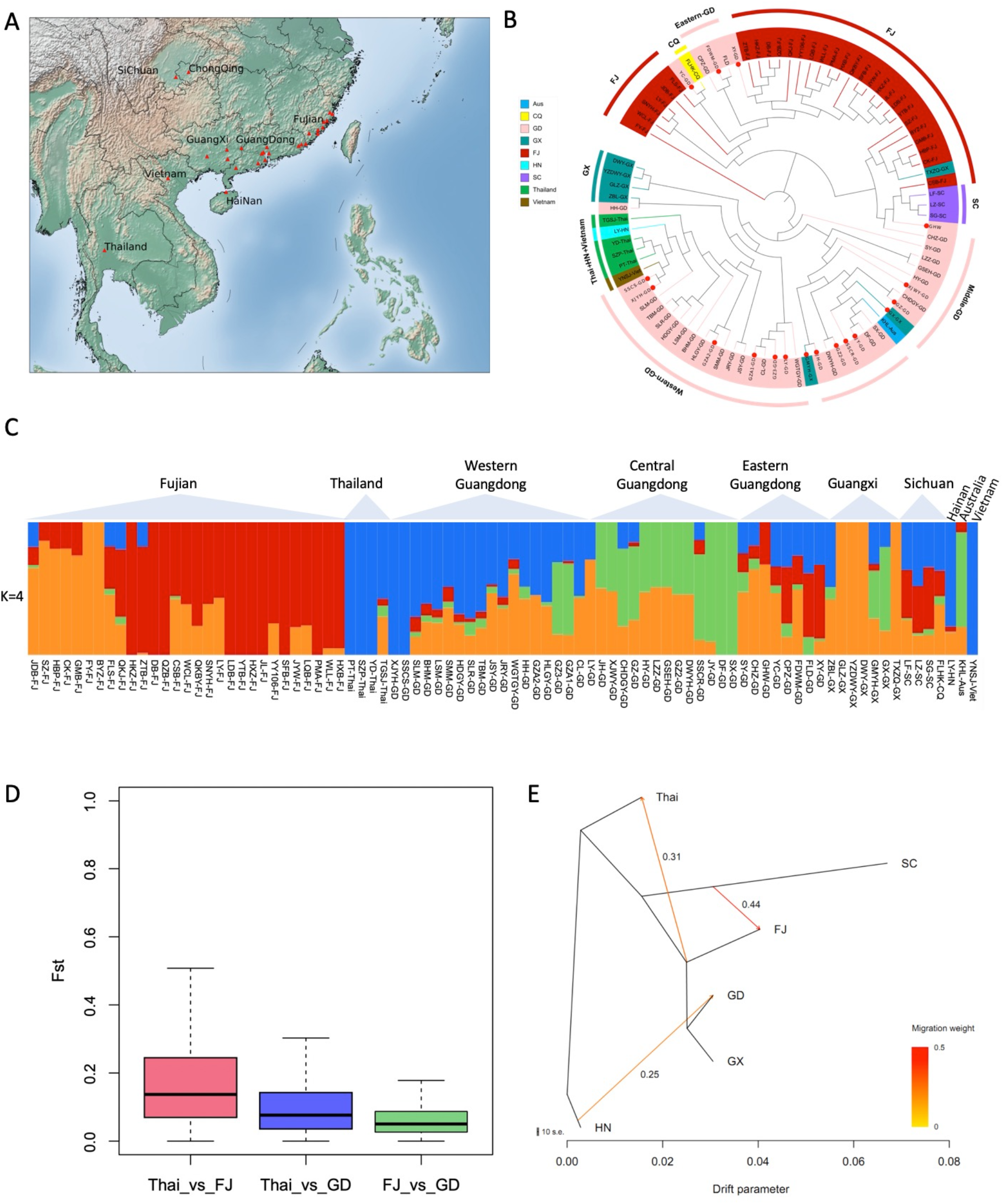
Population structure and admixture analysis of *Dimocarpus longan*. (**A**). Sampling locations of seven populations of *D. longan*, where red triangles distinguish the sampling locations (see Supplementary Table 9 for a complete list of samples). **(B)**. A neighbor-joining phylogenetic tree of all individuals of *D. longan* was constructed using SNPs. The artificial breeding individual was marked with red dots inside. Colors represent different geographic groups. **(C)**. A biogeographical ancestry (admixture) analysis of *D. longan* accessions with four ancestral clusters colored differently in the heatmap, where each column represents a longan sample. (**D)**. Distribution of *Fst* values (a measure of genetic differentiation) between longan population from Thailand (Thai), Fujian (FJ) and Guangdong (GD). **(E)**. maximum-likelihood tree and migration events among seven groups of *D. longan*. The migration events are colored according to their weight.

Phylogenomic analysis clustered 87 longan accessions into relatively distinct domestic Guangdong and Fujian groups after removal of artificial breeding populations (Figure 4B). Three Sichuan cultivars were next to Fujian group and distant to Guangdong group. Notably, two Guangdong cultivars, FLD and CPZ, were clustered with Fujian group, probably because they come from eastern Guangdong adjacent to Fujian. Guangdong cultivars are divided into two subgroups as “Shixai” (SX) centered group and “Chuliang” (CL) centered group from central and western Guangdong, respectively (Figure 4B), also the two main cultivars widely grown in Guangdong and Guangxi. Consistent with the phylogenetic tree, the principal component analysis showed that Guangdong and Fujian cultivars were overall grouped separately, while Thailand and Vietnam populations were distant to Chinese populations if artificial breeding cultivars were removed (Supplementary Figure 5).

To investigate the genetic background of longan from various regions, we performed biogeographical ancestry (admixture) analysis based on high-quality SNPs and tested it with ancestral group value (k) ranging from 2 to 8. With a choice of four ancestral groups (k=4) giving the smallest cross-validation errors (Supplementary Figure 6), the admixture analysis discovered a distinct genetic structure within longan accessions of different geographical origins. Longan cultivars from Fujian are composed of primarily two ancestral groups, whereas Guangdong, Guangxi and Sichuan cultivars contain fractions of all four ancestral groups, indicating their more complex ancestry backgrounds than Fujian ones (Figure 4C). The more similar ancestry composition between eastern Guangdong and Fujian cultivars is accordant to the geographical closeness of the two growing regions, suggesting their common ancestral origin or a possible exchange of cultivars between the two regions. By contrast, Thailand and Vietnam cultivars overall have a simple composition with predominantly one ancestral group, most likely shared with western Guangdong (Figure 4C). Thailand cultivars were genetically more related to western Guangdong cultivars (Figure 4B), but distant from Fujian cultivars. Consistent with this, we have also detected a stronger genetic differentiation (measured in *Fst* value) between Thailand and Fujian than between Thailand and Guangdong (Figure 4D). Notably, the Australian cultivar has a genetic background resembling middle Guangdong cultivar, suggesting it is a possible cultivar of middle-Guangdong origin introduced into Australia lately.

With the diverse ancestry backgrounds in these longan cultivars, we are curious about the migration history of longan germplasms and therefore investigated potential gene flows among different growing areas due to such migration using Treemix analysis. Given its reported origin in China, lots of wild longan resources are present in Yunnan and Hainan province of China^6^. Therefore, the Hainan cultivar was used as an outgroup in this analysis. The Treemix analysis detected a migration event directed from Hainan to Guangdong. The highest gene flow (migration weight 0.44) was observed between Sichuan and Fujian (Figure 4E). Gene flows were also detected from the Fujian, Guangdong and Guangxi populations to Thailand (migration weight 0.31) (Figure 4E). The detection of gene flows was consistent with longan migration history record. Longan was first cultivated in ‘Ling-nan’ district of China including Guangdong, Guangxi and Hainan about 2000 years ago, recorded by painting of “San Fu Huang”. According to history records, longan was moved to northern China-Shaanxi Province unsuccessfully, but was successfully introduced to Sichuan and then Fujian with suitable climate conditions (Yang Fu, “Chronicles of the South”, 1st century A.D.). Taken together, our analysis results overall matched history records that there was gene flow from Hainan wild germplasms to Guangdong, then a strong flow from Sichuan to Fujian, and finally the gene flow from China to Thailand.

### GWAS for longan fruit quality traits

We next performed GWAS for fruit qualities to uncover QTLs underlying these traits. Phenotyping data including pericarp thickness, pulp thickness, fruit horizontal diameter, total soluble solid, edible rate, and seed weight were collected from 80 longan cultivars from Longan Germplasm Repository of Guangdong Province, that were genotyped in this study. To validate the reliability of phenotype data, we calculated the Pearson Correlation Coefficient. Consistent with a previous study of longan^26^, the coefficient of variation of single fruit weight and seed weight was the largest, while the coefficient of variation of fruit weight, pericarp thickness and edible rate was the smallest (Supplementary Figure 7). GWAS analysis using 11,421,213 biallelic SNPs identified ten candidate QTLs significantly associated with six longan traits (p-value < 1e-6) (Supplementary Table 11), associating 12503, 7834, 11184, 6348, 46182 and 23523 SNPs with pericarp thickness, pulp thickness, fruit horizontal diameter, edible rate, seed weight and total soluble solid, respectively (Supplementary Figure 8). Genomic regions associated with seed weight contains several genes (Supplementary Table 9) related to seed development such as *UBQ10, UBIQ1, THI4, CBL10* and *CTPS2*. Specifically, a genomic region 0.94– 0.99 Mb on Chr6 contains three tandemly duplicated genes *DlBGL42* encoding β–glucosidase (Supplementary Table 12), consistent with reports that the seed size was controlled by *Clbg1* (watermelon β–glucosidase) due to decreased ABA content^27^.

The fruit sweetness is generally measured by total soluble solid content^28^. For a long time, high total soluble solid and edible rate are key longan traits selected for longan breeding^14^. Interestingly, GWAS identified a genomic region located at Chr3 (at ∼18.37 – 18.45 Mb) harboring QTLs shared between total soluble solid and seed weight (Figure 5A). A SNP variance analysis of this region found polymorphic site mutations of ∼37 kb at approximately 18.379 to 18.416 Mb, harboring three successive genes *DlPP304* (*D. long019821*.*01*), *DlACD6* (*D. long019822*.*01*) and *DlRDM3* (*D. long019823*.*01*) encoding mitochondria pentatricopeptide repeat-containing protein, accelerated cell death protein and an RNA-directed DNA methylation protein, respectively (Figure 5B). The co-localization of traits linked locus usually reflected positive significant correlation among different traits^29^. Sugar content is a characteristic of fruits that varies among cultivars and contributes to the distinctive flavor profile of longan varieties. There are reports that sugar contents are affected by fruit size and seed weight in sapodilla^30^, and soluble sugar accumulation affects Arabidopsis seed size^31^.

**Figure 5.**
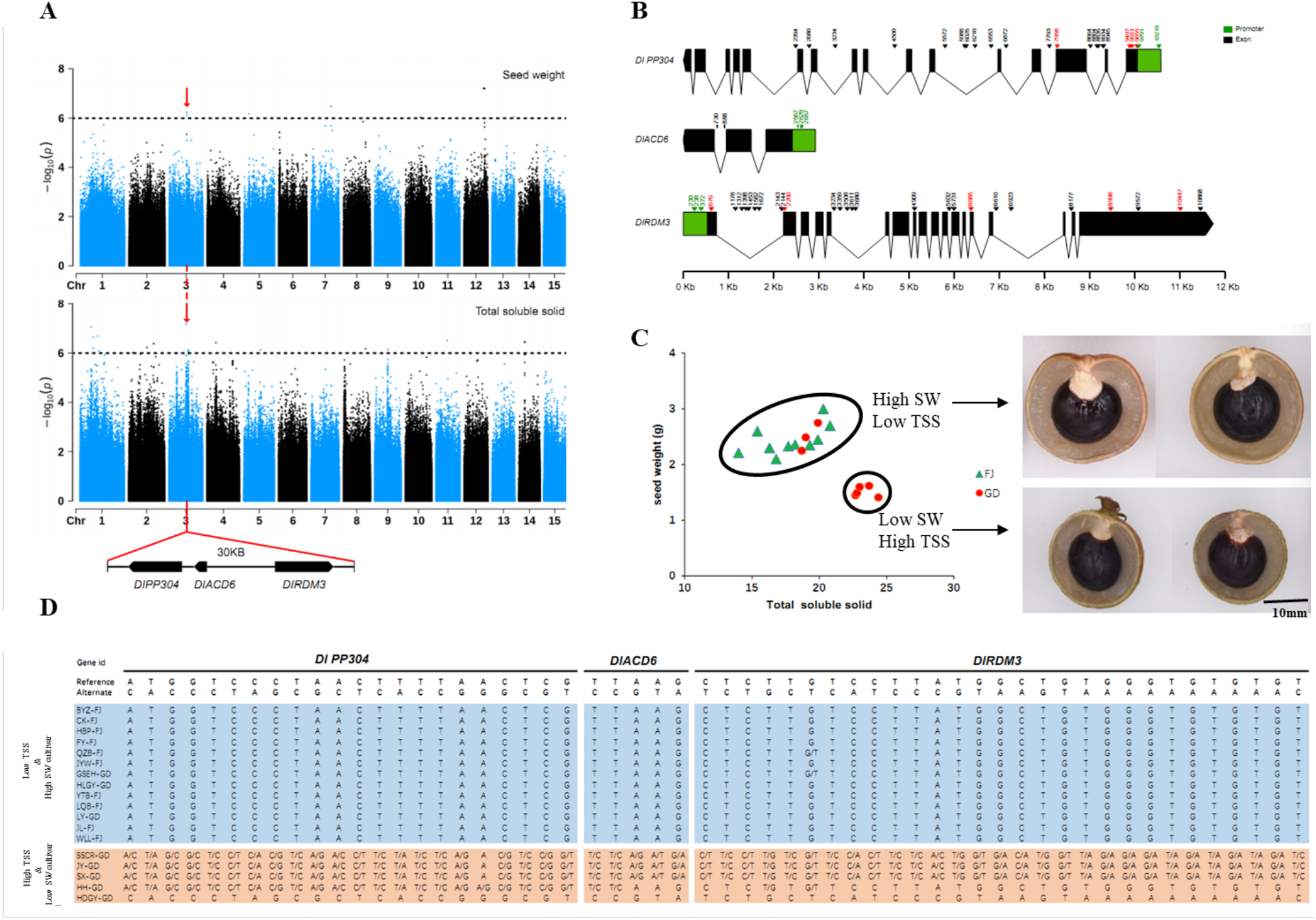
GWAS mapping of seed weight and total soluble solid in longan fruit. (**A**) Top: Manhattan plot summarizing the GWAS results for seed weight, based on analysis conducted with the randomly down sampled SNP call set. Middle: Manhattan plot of GWAS results for total soluble solid. Dotted lines represent the Bonferroni significance threshold. Red vertical lines represent overlapping region of two traits as highlighted below. Bottom: close-up of highlighted regions with three genes in the vicinity of the GWAS peak; (**B**) Diagram of three genes with respect to reference sequences and the haplotypes observed in samples collected from China. Black solid arrows indicate synonymous SNPs, Red solid arrows indicate non-synonymous SNPs specific to the high TSS (total soluble solid) with low SW (seed weight) varieties; (**C**) correlations of seed weight and TSS; (**D**) SNPs within the three genes for the most extreme TSS and SW phenotypes at each end of the high-type and low-type distributions (orange mutation is heterozygous to homozygous; blue: non-mutated) (see Supplementary Table 13 for all samples).

We then analyzed the seed weight and total soluble solid phenotypes in the 80 longan cultivars (Supplementary Figure 9), and observed that a group of ten Fujian and three Guangdong varieties displayed low soluble solid but with bigger and heavier seeds, whereas another group of five Guangdong varieties had high low soluble solid but smaller and lighter seeds (Figure 5C; Supplementary Figure 10). Examination of SNPs in the *DlPP304, DlACD6* and *DlRDM3* regions suggests that varieties of high soluble solid and low seed weight carried much more substantial number of variants than those of low soluble solid and high seed weight did (Figure 5D, Supplementary Table 11). Particularly, we found that the HDGY-GD variety carrying mostly homozygous variants at *DlPP304, DlACD6* and *DlRDM3* gene regions had the highest total soluble solid and lowest seed weight among the cultivars (Figure 5D). In summary, our GWAS analysis unveiled a list of QTLs and candidate genes for further validation using functional genomics or forward genetics approaches.

## DISCUSSION

Longan is one of the most important economic fruit trees in China and Southeast Asia. However, molecular breeding of longan cultivars is slow due to lack of genomic and biotechnological resources. Using a complement of single-molecule (PacBio) and Hi-C sequencing data, we have successfully produced a chromosome-level longan genome assembly followed by protein-coding gene annotations. This genome assembly for longan represents a significant improvement over a current draft reference genome assembly based on short-read sequencing data^9^. The improvement mainly stems from assembly of contigs using the PacBio long reads, followed by scaffolding based on chromatin interaction information given by Hi-C data. As primarily an outbreeding tree species, longan typically has high heterozygosity rate which presents challenge to contiguous assembly of its diploid genome. The JDB cultivar used for *de novo* genome assembly has a relatively low heterozygosity rate of 0.3% likely due to repeated inbreeding. Together the two factors facilitated the highly contiguous assembly of the genome. The repeat content (41%) in our genome assembly is less than what was reported previously for the assembly (52%) based on Illumina short reads by Lin *et al*.^9^, indicating that long read sequencing has helped resolved the ambiguous placement and assembly of repeat regions that were typically prone to mis-assembled by short reads. The high BUSCO score (98%) shows that completeness of longan assembly in our study is also improved compared to Lin *et al* assembly^9^, although we admit that there is still much to do for obtaining a complete longan genome sequence in the future. In short, we present a high-quality, chromosome-scale genome assembly of longan, the best of longan genome assemblies reported so far.

With the reference genome, we investigated the evolution of longan genome and showed that it shared the whole genome triplications and lacked additional WGDs. Lin *et al*. ^9^ performed 4DTv analysis their longan genome assembly to find that longan had an ancient genome duplication event, yet no information was given with respect to whether the event occurred in related plants. By contrast, we conducted inter-species analysis using longan and grape genome and showed that longan and grape had a 1:1 syntenic relations, confirming that the ancient duplication event in longan^9^ was actually a whole genome triplication (γ event) shared with grape and other core eudicots. Given that poplar genome underwent a post-γ WGD, the 1:1 and 1:2 syntenic relations between longan and grape, and between longan and poplar, suggested that the lack of WGD following the divergence of grape and longan, and of longan and poplar. Both Lin *et al*. ^9^ and our study identified a peak in *Ks* or 4DTv distribution for longan paralogs, indicating that many rounds of gene duplications occurred recently in longan genome, likely resulting from the activities of transposable elements.

The delightful taste and rich nutrition of longan fruits are among many unique traits of the plant. The longan is rich in polyphenolic compound in its fruits. Genomic analysis of longan can reveal critical information about how the plant evolved such special chemical reservoir. We detected significant gene family expansions related to biosynthetic genes of phenylpropanoids, terpenoids (sesquiterpenoid and triterpenoid) etc. compared to related plant lineages, suggesting that the longan genome has evolved towards innovation of these secondary metabolites due to natural or artificial selections. By contrast, Lin *et al*. ^9^ analyzed their draft longan genome assembly and also showed the longan genome experienced gene family expansions and contractions. However, no functional terms related to specialized metabolism were found to enrich in the expanded or contracted gene families they identified, although they showed enrichment of terms related to general biological processes such as cellular component organization etc. The different enrichment results may be down to several factors, such as an improved genome assembly and annotations, and selection of plant genomes for comparisons, and functional enrichment analysis tools etc. Our analysis also identified the expansion of UGTs gene families, which are essential for the diversification of structural features, and control the final oxidation, hydroxylation, and glycosylation steps to yield secondary metabolites in longan. The exact roles of UGTs in the formation and modification of specific longan natural products to give unique taste and nutrition of longan fruits remain to be determined.

Longan is widely grown in China and many Southeast Asian countries where longan cultivars were initially introduced from China, the origin of longan with the largest cultivation area in the world. Chinese longan mainly consists of Guangdong population, Fujian population and Guangxi population, and so on. Exchange of longan germplasms among different growing regions likely led to genetic introgression. Considering longan can undergo both inbreeding and outbreeding in natural environment, the genetic background and population structure of longan is rather complex and awaits elucidation. The 87 cultivated varieties in this study were mostly collected from mainly China and a few neighboring South Asian countries, which have excellent characteristics and are often used as breeding parents, but most evolved through long-term natural hybridization. The evolution model of natural hybridization is unclear, resulting in the unclear genetic background of germplasm resources. As a result, it is not uncommon to have confused names of longan varieties such as different varieties with the same name or same variety with the different names. Combining a high-quality genome assembly and population genomic sequencing of longan varieties, we have resolved the complex genetic background of longan, revealed its population admixture model and deduced the possible migration routes of longan in accordance with historic records. The high-quality SNP molecular markers generated in this study have enabled discovery of multiple QTLs strongly associated with useful traits such as seed weight and soluble solid, providing excellent resources for future molecular breeding. Taken together, these findings paved the way for future research through the integration of genomic, transcriptomic, and metabolomic data provide abundant promising candidates for phenylpropanoids biosynthesis.

## MATERIALS AND METHODS

### Germplasm genetic resources

A 30-year-old *D. longan* tree cultivar named JDB from the Longan Germplasm Repository of Guangdong Province (belonging to Institute of Fruit Tree Research at Guangdong Academy of Agricultural Sciences in China) was used for genome sequencing and *de novo* assembly in this study. Eighty-six additional *D. longan* cultivars (Supplementary Table 9) from the Longan Germplasm Repository of Guangdong Province, were collected for genome resequencing.

### DNA and RNA isolation

Longan cultivar JDB was planted in Longan Germplasm Repository of Guangdong Province. The fresh and healthy young leaves were collected, cleaned and used for genomic DNA isolation and sequencing. Genomic DNA was extracted from young fresh leaves of *D. longan* using the modified cetyltrimethylammonium bromide (CTAB) method^32^. The concentration and purity of the extracted DNA were assessed using a Nanodrop 2000 spectrophotometer (Thermo, MA, USA) and Qubit 3.0 (Thermo, CA, USA), and the integrity of the DNA was measured using pulsed-field electrophoresis with 0.8% agarose gel. In addition, fresh leaves and other tissues (roots, shoots, young fruits) of JDB cultivar were collected for RNA isolation and transcriptome sequencing. Total RNA was isolated with RNAprep Pure Plant Kit (Tiangen Biotech) according to the manufacturer’s instructions. The integrity and quantity of extracted RNA were analyzed on an Agilent 2100 Bioanalyzer. For each tissue, three biological replicates were prepared for sequencing.

### Genome and transcriptome sequencing

DNA sequencing libraries were constructed and sequenced on the Illumina NovaSeq 6000 platform at 50x depth according to the manufacturer’s protocols (Illumina). To generate long-read sequencing reads for *D. longan*, DNA libraries for PacBio SMRT sequencing were prepared following the PacBio standard protocols and sequenced on a Sequel platform. In brief, genomic DNA was randomly sheared to an average size of 20 kb, using a g-Tube (Covaris). The sheared gDNA was end-repaired using polishing enzymes. After purification, a 20-kb insert SMRTbell library was constructed according to the PacBio standard protocol with the BluePippin size-selection system (Sage Science) and sequences were generated on a PacBio Sequel (9 cells) and PacBio RS II (1 cell) platform by Biomarker Technologies. Raw subreads was filtered based on read quality (≥0.8) and read length (≥1000bp). For chromosome-level genome scaffolding, Hi-C libraries were prepared from fresh leaves and sequenced on the Illumina HiSeq X Ten platform. DNA was digested with HindIII enzyme, and the ligated DNA was sheared into size of 200-400bp. The resulting libraries was sequenced by using Illumina NovaSeq 6000. For transcriptome sequencing, RNA sequencing (RNA-seq) libraries were constructed using True-Seq kit (Illumina, CA), and sequenced using Illumina HiSeq X Ten platform. Illumina raw reads were trimmed using *Trimmomatic* (v0.39)^33^ with parameters “LEADING: 10 TRAILING:10 SLIDINGWINDOW:3:20 MINLEN:36” to remove adapter sequences and low-quality reads, yielding a total of ∼77.7 Gb clean RNA-seq data from four tissues.

### Genome assembly and evaluation

To estimate the genome size and heterozygosity level of *D. longan*, cleaned Illumina PE reads were used for k-mer spectrum analysis using *GenomeScope* (v2.0)^34^ based on 21-mer statistics. PacBio SMRT reads were used for de novo genome assembly by using *Canu* (V1.9)^15^ pipeline with parameters “correctedErrorRate=0.045 corMhapSensitivity=normal ‘batOptions=-dg 3 -db 3 -dr 1 -ca 500 -cp 50”. Alternative haplotig sequences was removed using *purge_dups*^35^ according default settings, and only primary contigs were kept for downstream analysis. To correct the base-pair level errors in raw assembly sequences, two rounds of polishing were conducted using high-qualitied Illumina DNA reads with *Pilon* (v1.23)^16^. The longan contigs were further anchored to chromosomes using *3D-DNA*^17^ based on Hi-C contact map, followed by manual correction using *Juicerbox* (v1.11.08)^36^ to fix assembly errors. The completeness of genome assembly was assessed by BUSCO v1.22^18^ using 2121 eudicotyledons_odb10 single copy genes. PacBio sequence reads and Illumina DNA reads were aligned to the genome sequences using *minimap2*^37^ and BWA^38^ respectively.

### Repetitive element annotation

We used a combination of the *de novo* repeat library and homology-based strategies to identify repeat structures. *TransposonPSI*^39^ was used to identify transposable elements. *GenomeTools* suite^40^ (LTR harvest and LTR digest) was used to annotate LTR-RTs with protein HMMs from the Pfam database. Then, a *de novo* repeat library of longan genome was built using *RepeatModeler*^41^, and each of the three repeat libraries was classified with *Repeat_Classifier*. Subsequently, the non-redundant repeat library was analyzed using BLASTx to search the transposase database (evalue=1e-10) and non-redundant plant protein databases (evalue=1e-10) to remove protein-coding genes. Then, the *de novo* repeat library was used to discover and mask the assembled genome with *RepeatMasker*^42^ with the “-xsmall -excln” parameter.

### Prediction and annotation of protein-coding genes

For gene structure annotations, the RNA-seq data of four different tissues were aligned to repeat-soft masked genome using *STAR*^43^, which generates intron hints for gene structure annotation. The structural annotation of protein-coding genes was performed using *BRAKER2*^44^ by combining the aligned results from *ab initio* predictions, homologous protein mapping, and RNA-seq mapping to produce the final gene prediction. The genes with protein length < 120 amino acids and expression level < 0.5 TPM were removed. Predicted genes were assigned functions by performing BLAST against the NCBI non-redundant protein database with e-value threshold of 1e-10. In addition, a comprehensive annotation was also performed using *InterProScan* (5.36-75.0)^45^.

### Comparative genomics analysis

Putative orthologship was constructed using protein sequences from two monocots, ten eudicots and *Amborella trichopoda* and longan. Only longest protein sequence was selected as representative of each gene. Orthogroups were inferred by *OrthoFinder* (v2.4.1)^46^. The species tree was used as a starting tree to estimate species divergence time using *MCMCTREE* in *paml* (v4.9)^47^ package. Speciation event dates for *Ananas comosus*-*Oryza sativa* (102∼120 MYA), *Populus trichocarpa*-*Ricinus communis* (70∼86 MYA), *Arabidopsis thaliana*-*Carica Papaya* (63∼82 MYA), and *Glycine max*-*Citrus sinensis* (98∼117 MYA) obtained from *Timetree* (www.timetree.org) were used to calibrate the divergence time estimation. We conducted two independent *MCMCTREE* runs using the following settings: burnin = 20000, sampfreq = 30, and nsample = 20000.

The ortholog count table and phylogenetic tree topology inferred from the *OrthoFinder* were provided to *CAFÉ* (v4.2)^48^, which identifies significant expansion or contraction occurred in each gene family across species using a random birth and death model to estimate the size of each family at each ancestral node. Among expanded gene families, longan genes enriched with IPR002213 (UDP-glucuronosyl/UDP-glucosyltransferase) and IPR036396 (Cytochrome P450 superfamily) and their ortholog CDS sequences of *A. thaliana* and *C. sinensis* genome were retrieved. Multiple sequence alignment was conducted using *MUSCLE* (v3.8.1551)^49^ software. *IQtree* was used to constructed a maximum likelihood tree with parameters “-m MF”. Tree file was loaded into the Interactive Tree of Life (iTOL) web server for tree visualization and figure generation^50^.

### Transcriptomic analysis

After removing adapters and trimming low-quality bases, RNA-seq reads were mapped to the longan reference genome using *STAR*^43^ with parameters “--alignIntronMax 6000 –align IntronMin 50” and then using *RSEM* tool^51^ for transcripts quantification. Outliers among the individual experimental samples were verified based on the Person correlation coefficient, *r*^2^ ≥0.85. Differential expression analysis was performed using *DEseq2*^52^ package. Genes were differentially expressed between two conditions if the adjusted p-values was <0.01 and fold change > 1.

### Genetic variation detection

Population resequencing reads were mapped to the chromosome-level genome assembly of longan using *BWA*^*38*^ with default parameters. Alignments for each sample were processed by removing duplicate reads using the *Samtools*^53^. The mpileup function in *Samtools* was used to generate mpileup files for each sample. *Bcftools*^54^ was used to identify SNPs and small indels. The SNPs were filtered with a criterium of read mapping quality (MQ) higher than 40, minimum coverage greater than 10, base quality more than 30, and genotype missing rate < 20% (of all samples). The high quality SNPs were used in further analysis.

### Population structure analysis

The vcf-format SNP sets were transformed into binary ped-format using *PLINK*^55^. To estimate individual admixture assuming different numbers of clusters, the population structure and ancestry were investigated using *ADMIXTURE*^56^ based on all SNPs. A linkage disequilibrium pruning step was performed with *PLINK*^55^ with parameters “--indep-pairwise 50 10 0.1”. We analyzed the number (K) of ancestry clusters ranging from 2 to 8, and found that the cross validation error was smallest at K=4. We also inferred a population-level phylogeny across all groups, using the maximum likelihood approach implemented in *TreeMix* ^57^. For such analysis, the samples with the genotype missing rate > 20% were filtered out. The SNPs with an imputation info score of < 0.8, minor allele frequency (MAF) < 5%, and significant deviation (p < 10e-4) from Hardy-Weinberg equilibrium (HWE) also were removed.

### GWAS analysis

Phenotypic data of the *D. longan* cultivars in Longan Germplasm Repository of Guangdong Province, including pericarp thickness, pulp thickness, fruit horizontal diameter, total soluble solid, edible rate, and seed weight, were measured in 2020 and 2021. Fifty mature fruits per cultivar were harvested for phenotypic measurement from five longan trees of forty years old, which had even canopy size and fruit load. To ensure the statistical power of analysis, we used the SNPs of MAF > 5% to represent the genotyping information for each GWAS. Gamma (version 0.98.1) was utilized to carry out the GWAS analyses with a mixed linear modal (MLM)^58^. Furthermore, to minimize false positives in GWAS, population structure was taken into account using a kinship matrix that was estimated using *PLINK*^55^. The genome-wide significance threshold was set at 1e-6.

## Data availability

The genome sequencing data, RNA-seq data, Hi-C data, genome assembly of *D. longan* generated in this study have been deposited in the Genome Sequence Archive (GSA) database at National Genomics Data Center of China National Center for Bioinformation under accession number CRA004281.

## AUTHOR CONTRIBUTIONS

Project design and oversight: LG, JL and WQ; Sample collection and curation: DG and SH; Conducting experiment and data analysis: JW, ZL and LG; Result interpretation: LG, JW, JL, BL and WQ; Figure and table preparation: LG, JW and LZ; Manuscript writing and revision: LG, JW, BL and WQ; Provide funding: JL and LG; All authors have read and proved the final version of this manuscript.

## ACKNOWLEDGEMENT

This project is supported by Key-Area Research and Development Program of Guangdong Province (2020B020220006) and Guangdong Provincial Crops Germplasm Nursery Construction and Resources Collection, Preservation, Identification & Evaluation Foundation. In addition, LG is supported by the National Natural Science Foundation of China (31970317) and faculty startup package from Peking University Institute of Advanced Agriculture Sciences. The authors also would like to thank anonymous reviewers for their comments and suggestions to improve this manuscript.

## SUPPLEMENTARY MATERIALS

Supplementary Figure 1: The heatmap of phenylalanine ammonia-lyase genes (PALs) expressed in various longan tissues.

Supplementary Figure 2: The heatmap of peroxidase genes (PODs) expressed in various *Dimocarpus longan* tissues.

Supplementary Figure 3: InterPro protein domain enrichment analysis of *Dimocarpus longan* expanded gene families.

Supplementary Figure 4: The heatmap of UGTs genes expressed in various longan tissues.

Supplementary Figure 5: Principal component analysis of *Dimocarpus longan* samples based on genotypes.

Supplementary Figure 6: Biogeographical ancestry analysis with group value K.

Supplementary Figure 7: Pearson correlation coefficient matrix for analyses of quantitative traits related to fruit quality in longan germplasms.

Supplementary Figure 8: Manhattan plot for the genome wide association analysis of longan fruit traits. Supplementary Figure 9: Phenotypes of seed and TSS.

Supplementary Table 1: Sequencing statistics.

Supplementary Table 2: Summary of Illumina data for genome survey and genome polishing.

Supplementary Table 3: Gene function annotated by different databases.

Supplementary Table 4: Statistics of repetitive elements.

Supplementary Table 5: Comparison of genes in orthogroups between *Dimocarpus longan* and 13 other species.

Supplementary Table 6: List of expanded phenylpropanoid biosynthesis genes and their expression level in different tissues.

Supplementary Table 7: List of different expressed IPR enriched gene families.

Supplementary Table 8: List of UGTs genes ID and their expression level.

Supplementary Table 9: List of genome resequencing samples and their locations.

Supplementary Table 10: GWAS results of significant SNPs associated with six traits and their annotations.

Supplementary Table 11: SNPs at three genes related to TSS and seed weight.

