## Supplementary Figures for "Genomic insights into longan evolution from a chromosome-level genome assembly and population genomics of longan accessions"

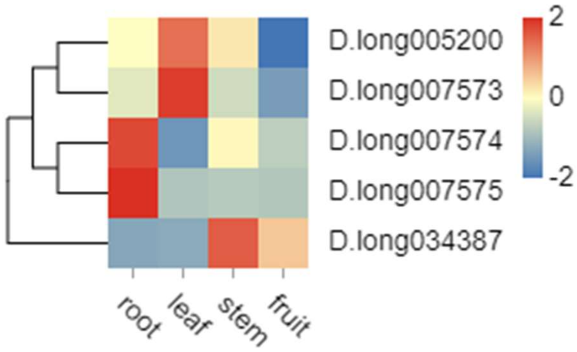


**Supplementary Figure 1: The heatmap of phenylalanine ammonia-lyase genes (PALs) expressed in various longan tissues.**

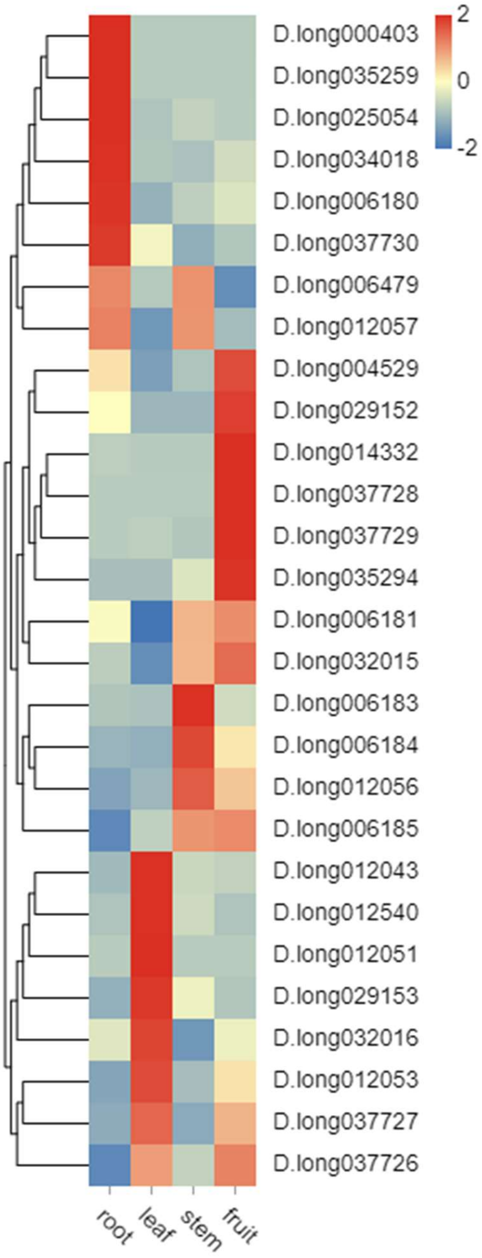


**Supplementary Figure 2: The heatmap of peroxidase genes (PODs) expressed in various *Dimocarpus longan* tissues.**

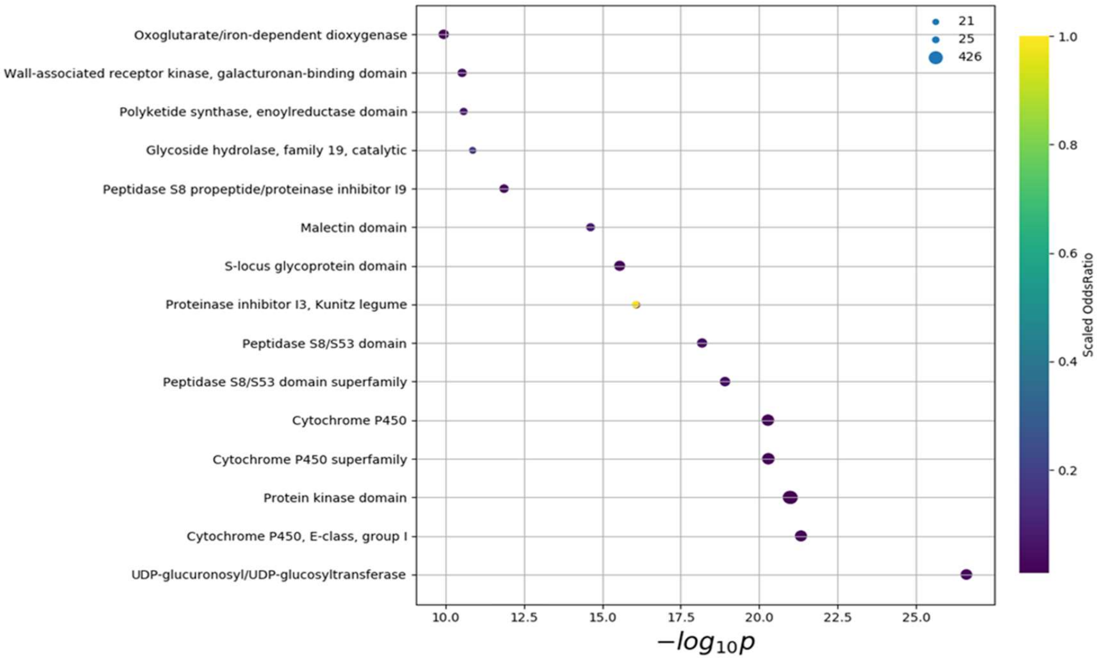


**Supplementary Figure 3: InterPro protein domain enrichment analysis of *Dimocarpus longan* expanded gene families.**

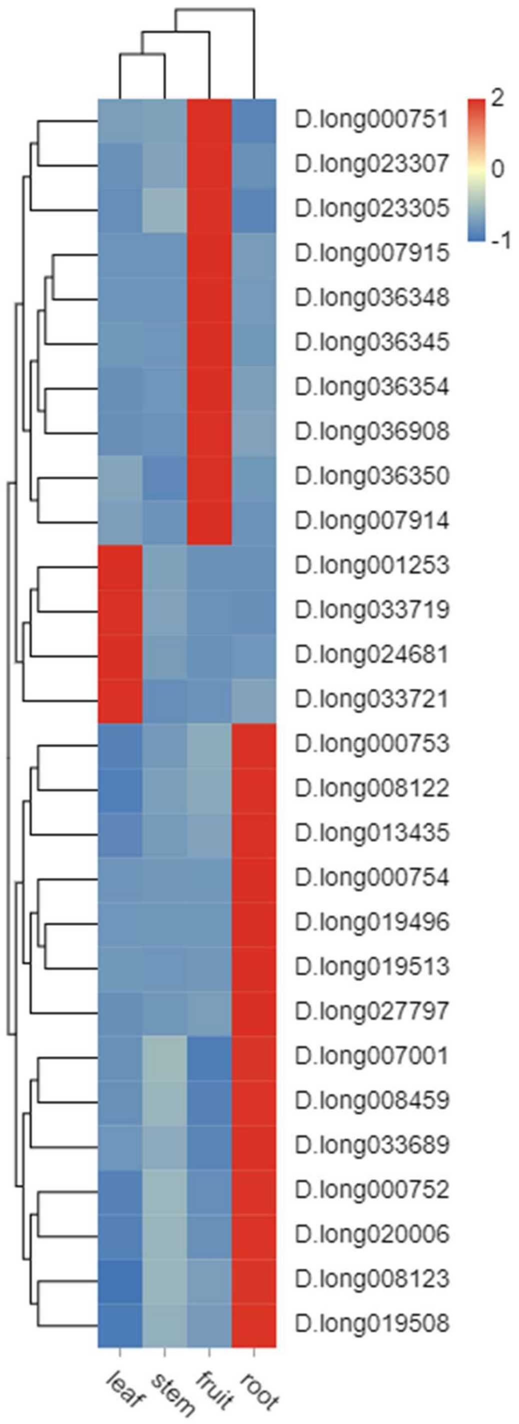


**Supplementary Figure 4: The heatmap of UGTs genes expressed in various longan tissues.**

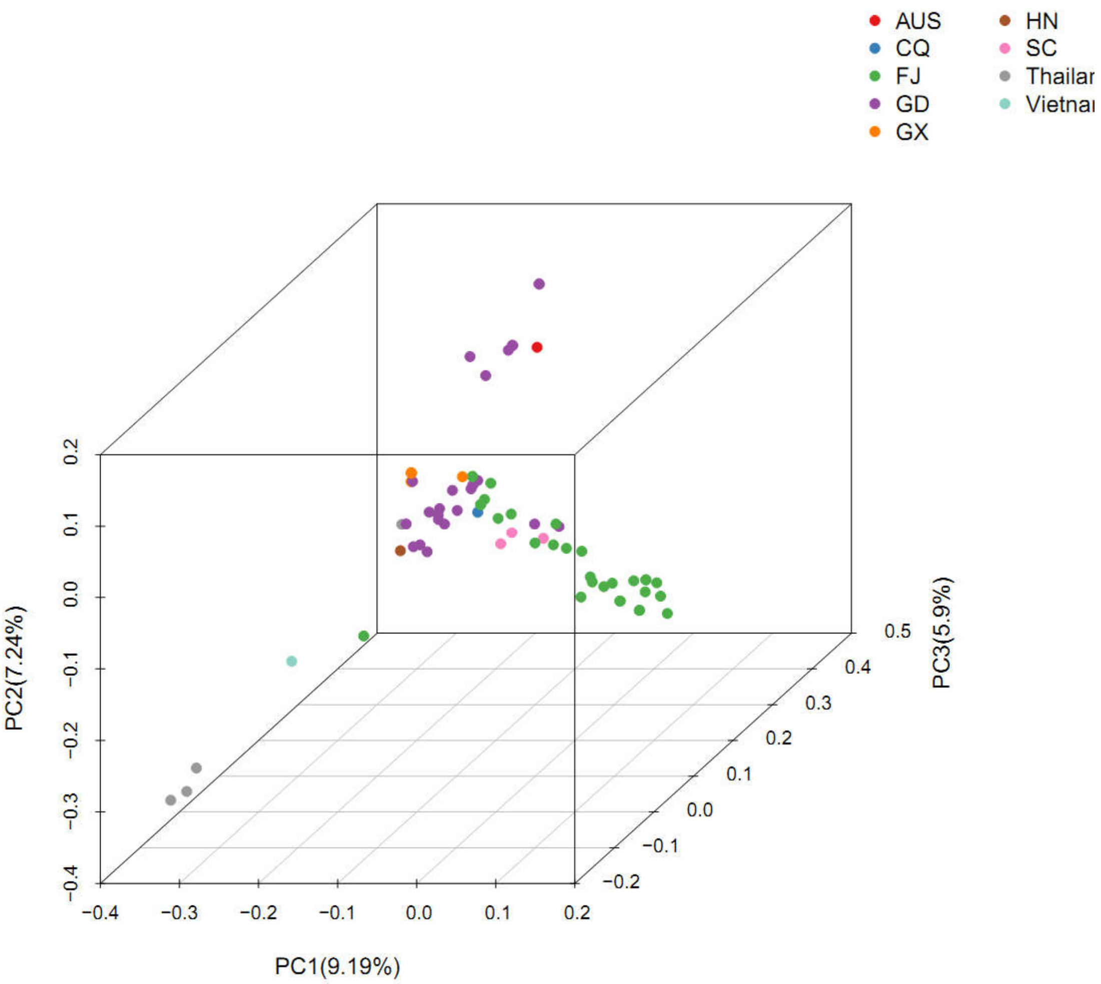


**Supplementary Figure 5: Principle component analysis of *Dimocarpus longan* samples based on genotypes.**

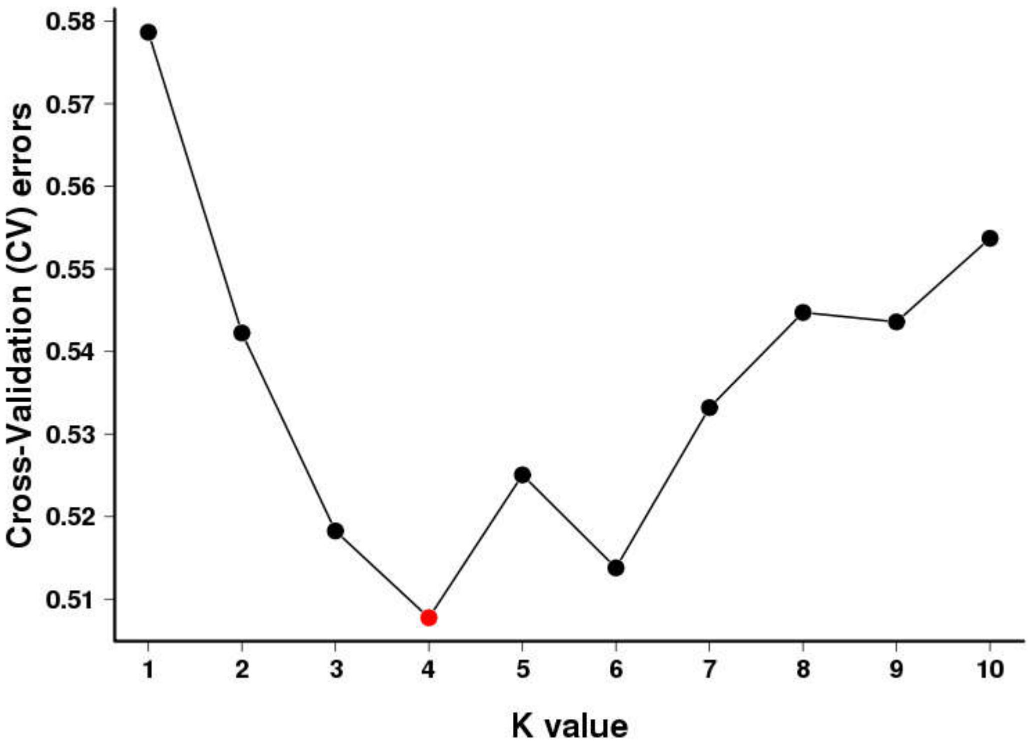


**Supplementary Figure 6: Biogeographical ancestry analysis with group value K.**

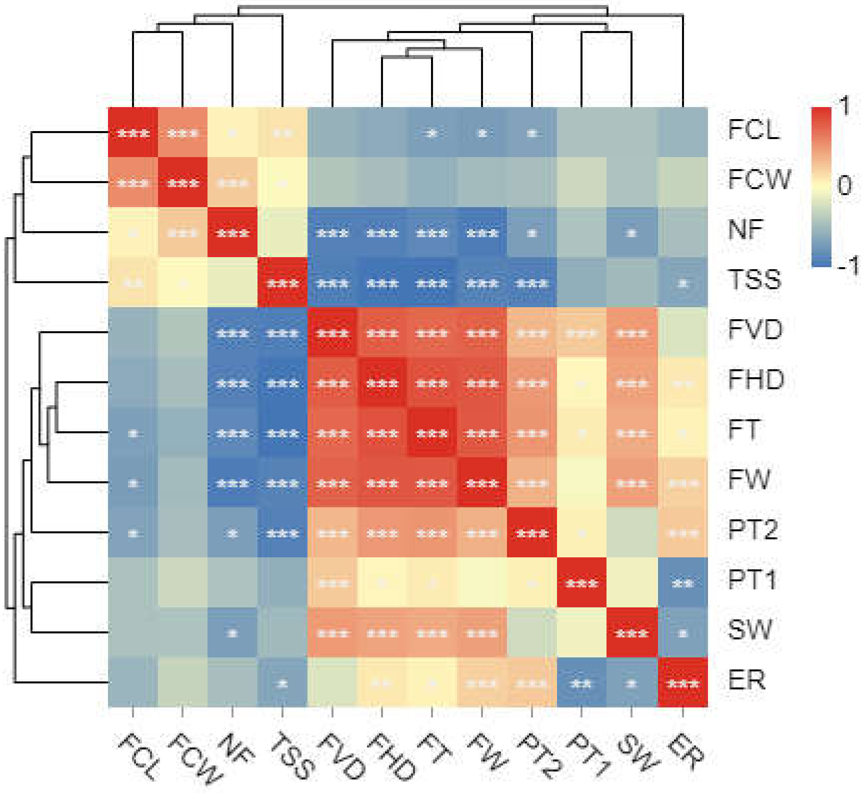


**Supplementary Figure 7: Pearson correlation coefficient matrix for analyses of quantitative traits related to fruit quality in 71 germplasms.**

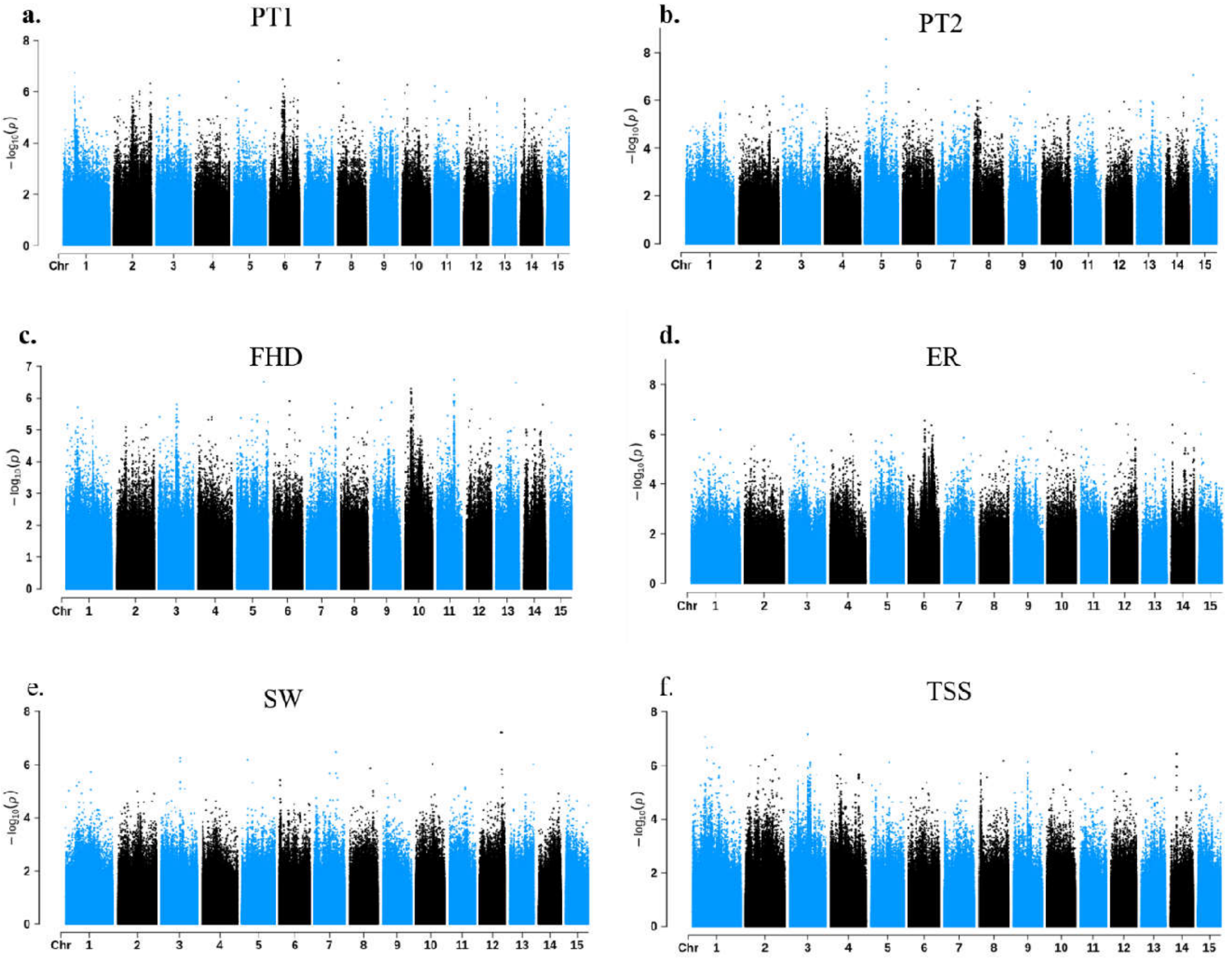


**Supplementary Figure 8: Manhattan plot for the genome wide association analysis of longan fruit traits.** (a) Fruits' pericarp thickness (PT1). (b) Fruits' pulp thickness (PT2). (c) Fruits' horizontal diameter (FHD). (d) Fruits' edible percentage (ER). (e) Seed weight (SW). (f) Total soluble solid (TSS).

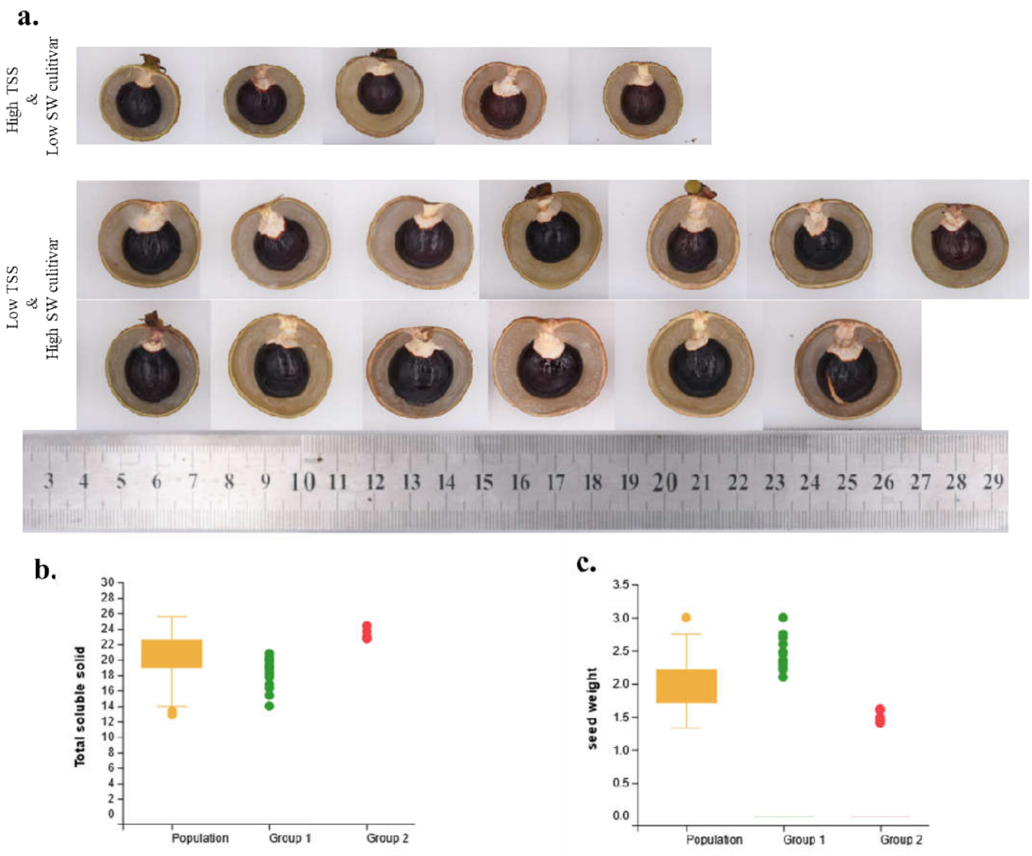


**Supplemental Figure 9: Phenotypes of seed and TSS.** (a) top: varieties with lower TSS and higher SW, bottom: varieties with higher TSS and lower SW; (b) distribution of TSS among cultivars with photos here; (c) distribution of seed weight among cultivars with photos here.
