## Supplementary Tables for "Genomic insights into longan evolution from a chromosome-level genome assembly and population genomics of longan accessions"

Table S1. Sequencing statistics.

| | | |
|---|---|---|
| **#Illumina NGS sequence** | | |
| **total base** | **readlen** | **coverage** |
| 25.3G | 150bp | 56X |
| **#PacBio** | | |
| **total base** | **Sequences No.** | **Sequences Max.** |
| 184.4G | 8452962 | 441148 |
| | Mean length | N50 |
| | 21819 | 32005 |
| **#HiC** | | |
| **total base** | **readlen** | **coverage** |
| 57.6G | 150bp | 127X |

Table S2. Summary of Illumina data for genome survey and genome polishing.

| | |
|---|---|
| **Sequences NO.:** | 250 |
| **Sequences Min.:** | 1,406 |
| **Sequences Max.:** | 31,067,541 |
| **Mean length:** | 1,821,787 |
| **N50:** | 12,096,926 |
| **Total number of bases:** | 455,446,992 |
| **GC%** | 34% |
| **Pacbio Reads mapping rate** | 89.92% |
| **Illumina Reads mapping rate** | 95.50% |
| **Genome coverage rate** | 94.60% |
| **BUSCO[genome mode]** | C:96.5%[S:92.6%,D:3.9%],F:1.6%,M:1.9%,n:2121 |
| **LAI index** | 20.78 |
| **Complete BUSCOs (C)** | 2048 |
| **Complete and single-copy BUSCOs (S)** | 1965 |
| **Complete and duplicated BUSCOs (D)** | 83 |
| **Fragmented BUSCOs (F)** | 33 |
| **Missing BUSCOs (M)** | 40 |
| **Total BUSCO groups searched #eudicotyledons_odb10** | 2121 |

Table S3. Gene function annotated by different databases.

| | **Annotated number** | **Percent of total genes** |
|---|---|---|
| **NR** | 35,959 | 89.00% |
| **KEGG** | 34,152 | 84.60% |
| **PlantTFDB** | 1,709 | 4% |
| **InterPro** | 29,661 | 73.40% |

Table S4. Statistics of repetitive elements.

| | | | Number | Length(bp) | % of whole genome |
|---|---|---|---|---|---|
| **Class I: Retroelement** | **LTR elements** | **Copia** | 54,443 | 38,850,596 | 8.552 |
| | | **Gypsy** | 72,339 | 70,559,178 | 15.533 |
| | | **other** | 10,533 | 5,561,251 | 1.224 |
| | **Non-LTR elements** | **LINE** | 24,872 | 8,649,078 | 1.904 |
| | | **SINE** | 78 | 11,196 | 0.002 |
| | | **other** | 1,324 | 367,787 | 0.081 |
| **Class II:DNA transposon** | **DNA/Helitron** | | 5,661 | 2,106,028 | 0.464 |
| | **DNA/Harbinger** | | 8,957 | 3,276,187 | 0.721 |
| | **DNA/hAT** | | 36,925 | 12,647,303 | 2.784 |
| | **DNA/Crypton** | | 1,043 | 241,125 | 0.053 |
| | **DNA/EnSpm** | | 23,604 | 8,370,445 | 1.843 |
| | **DNA/Mariner** | | 1,659 | 401,375 | 0.088 |
| | **DNA/MuDR** | | 25,436 | 11,839,905 | 2.606 |
| | **DNA/Polinton** | | 3,214 | 931,063 | 0.205 |
| | **DNA/other** | | 28,163 | 8,563,583 | 1.885 |
| **Tandem_Repeat** | | | 1,838 | 2,025,952 | 0.446 |
| **Unclassified elements** | | | 93,696 | 21,041,193 | 4.632 |
| | | | | 189,423,498 | 41.699 |

Table S5. Comparison of genes in orthogroups between Dimocarpus longan and 13 other species.

| | Amborella trichopod | Ananas_comosus | Arabidopsis_thaliana | Carica_Papaya | Citrus_sinensis | Dimocarpus_longan | Glycine_max | Nicotiana_attenuata | Oryza_sativa | Populus_trichocarpa | Ricinus_communis | Solanum_tuberosum | Theobroma_cacao | Vitis_vinifera |
|---|---|---|---|---|---|---|---|---|---|---|---|---|---|---|
| **Number of genes** | 27313 | 27024 | 27628 | 27751 | 25379 | 40420 | 55897 | 33320 | 35775 | 41335 | 31221 | 39021 | 21330 | 29927 |
| **Number of genes in orthogroups** | 22563 | 24366 | 25219 | 23072 | 23709 | 37401 | 49595 | 31860 | 27899 | 36440 | 23922 | 36101 | 20984 | 25725 |
| **Number of unassigned genes** | 4750 | 2658 | 2409 | 4679 | 1670 | 3019 | 6302 | 1460 | 7876 | 4895 | 7299 | 2920 | 346 | 4202 |
| **Percentage of genes in orthogroups** | 82.6 | 90.2 | 91.3 | 83.1 | 93.4 | 92.5 | 88.7 | 95.6 | 78 | 88.2 | 76.6 | 92.5 | 98.4 | 86 |
| **Percentage of unassigned genes** | 17.4 | 9.8 | 8.7 | 16.9 | 6.6 | 7.5 | 11.3 | 4.4 | 22 | 11.8 | 23.4 | 7.5 | 1.6 | 14 |
| **Number of orthogroups** | 13258 | 13108 | 12871 | 13480 | 13498 | 14712 | 15148 | 14262 | 13871 | 14466 | 14349 | 13979 | 13063 | 14107 |
| **Percentage of orthogroups containing species** | 45.2 | 44.6 | 43.8 | 45.9 | 46 | 50.1 | 51.6 | 48.6 | 47.2 | 49.3 | 48.9 | 47.6 | 44.5 | 48 |
| **Number of species-specific orthogroups** | 902 | 711 | 694 | 501 | 227 | 957 | 1588 | 616 | 1623 | 702 | 767 | 803 | 103 | 617 |
| **Number of genes in species-specific orthogroups** | 4485 | 4067 | 3252 | 2009 | 703 | 6510 | 6105 | 3481 | 6079 | 2757 | 2529 | 8138 | 435 | 2046 |
| **Percentage of genes in species-specific orthogroups** | 16.4 | 15 | 11.8 | 7.2 | 2.8 | 16.1 | 10.9 | 10.4 | 17 | 6.7 | 8.1 | 20.9 | 2 | 6.8 |

| **Number of species** | **Number of genes** | **Number of genes in orthogroups** | **Number of unassigned genes** | **Percentage of genes in orthogroups** | **Percentage of unassigned genes** | **Number of orthogroups** | **Number of species-specific orthogroups** | **Number of genes in species-specific orthogroups** | **Percentage of genes in species-specific orthogroups** | **Mean orthogroup size** | **Median orthogroup size** | **G50 (assigned genes)** | **G50 (all genes)** | **O50 (assigned genes)** | **O50 (all genes)** | **Number of orthogroups with all species present** | **Number of single-copy orthogroups** |
|---|---|---|---|---|---|---|---|---|---|---|---|---|---|---|---|---|---|
| 14 | 463341 | 408856 | 54485 | 88.2 | 11.8 | 29360 | 10811 | 52596 | 11.4 | 13.9 | 9 | 22 | 20 | 4901 | 6199 | 7530 | 137 |

Table S6. List of expanded phenylpropanoid biosynthesis genes and their expression level in different tissues.

| #ID | SEQ_Length | Database | Function_ID | Function_Name | root-average expression | leaf-average expression | stem-average expression | fruit-average expression |
|---|---|---|---|---|---|---|---|---|
| **D.long005483** | 306 | Pfam | PF08240 | Alcohol dehydrogenase GroES-like domain | 0 | 0 | 0 | 0 |
| **D.long024265** | 356 | Pfam | PF08240 | Alcohol dehydrogenase GroES-like domain | 0 | 0.06 | 0 | 0.026666667 |
| **D.long030408** | 355 | Pfam | PF08240 | Alcohol dehydrogenase GroES-like domain | 282.4866667 | 0.153333333 | 43.41666667 | 16.1 |
| **D.long030412** | 287 | Pfam | PF08240 | Alcohol dehydrogenase GroES-like domain | 32.70666667 | 1.476666667 | 77.27333333 | 26.30333333 |
| **D.long030414** | 354 | Pfam | PF08240 | Alcohol dehydrogenase GroES-like domain | 32.08333333 | 113.3433333 | 18.26 | 51.70333333 |
| **D.long030415** | 355 | Pfam | PF08240 | Alcohol dehydrogenase GroES-like domain | 203.8633333 | 3.86 | 33.23333333 | 22.76 |
| **D.long031881** | 362 | Pfam | PF08240 | Alcohol dehydrogenase GroES-like domain | 201.3966667 | 1.946666667 | 207.4433333 | 45.2 |
| **D.long031882** | 360 | Pfam | PF08240 | Alcohol dehydrogenase GroES-like domain | 0.653333333 | 0 | 0.116666667 | 6.78 |
| **D.long031883** | 362 | Pfam | PF08240 | Alcohol dehydrogenase GroES-like domain | 1.03 | 3.546666667 | 9.786666667 | 2.1 |
| **D.long031884** | 361 | Pfam | PF08240 | Alcohol dehydrogenase GroES-like domain | 1.543333333 | 0.21 | 1.436666667 | 3.733333333 |
| **D.long031885** | 346 | Pfam | PF08240 | Alcohol dehydrogenase GroES-like domain | 2.85 | 0 | 1.143333333 | 0.706666667 |
| **D.long031886** | 362 | Pfam | PF08240 | Alcohol dehydrogenase GroES-like domain | 19.00333333 | 0.243333333 | 3.29 | 7.24 |
| **D.long031887** | 362 | Pfam | PF08240 | Alcohol dehydrogenase GroES-like domain | 1.523333333 | 0.806666667 | 22.91666667 | 3.57 |
| **D.long031892** | 361 | Pfam | PF08240 | Alcohol dehydrogenase GroES-like domain | 86.49333333 | 0 | 4.226666667 | 0.153333333 |
| **D.long032679** | 360 | Pfam | PF08240 | Alcohol dehydrogenase GroES-like domain | 0 | 154.7566667 | 1.953333333 | 81.24 |
| **D.long032684** | 359 | Pfam | PF08240 | Alcohol dehydrogenase GroES-like domain | 5.373333333 | 5.913333333 | 16.55 | 16.65666667 |
| **D.long003186** | 501 | Pfam | PF00171 | Aldehyde dehydrogenase family | 0.156666667 | 2.696666667 | 0.873333333 | 0.303333333 |
| **D.long014098** | 501 | Pfam | PF00171 | Aldehyde dehydrogenase family | 1.046666667 | 105.1966667 | 19.39 | 146.3666667 |
| **D.long002765** | 548 | Pfam | PF00501 | AMP-binding enzyme | 96.25 | 55.25333333 | 86.36 | 36.34333333 |
| **D.long011121** | 540 | Pfam | PF00501 | AMP-binding enzyme | 6.646666667 | 159.22 | 119.8133333 | 63.55333333 |
| **D.long011346** | 540 | Pfam | PF00501 | AMP-binding enzyme | 211.4266667 | 191.85 | 139.7733333 | 78.69333333 |
| **D.long014725** | 580 | Pfam | PF00501 | AMP-binding enzyme | 13.83 | 29.06 | 74.93 | 76.85333333 |
| **D.long024147** | 493 | TIGRFAM | TIGR03356 | BGL: beta-galactosidase | 74.56 | 90.94333333 | 128.1166667 | 125 |
| **D.long024148** | 493 | TIGRFAM | TIGR03356 | BGL: beta-galactosidase | 100.47 | 18.82333333 | 19.98333333 | 56.21666667 |
| **D.long024149** | 493 | TIGRFAM | TIGR03356 | BGL: beta-galactosidase | 183.3133333 | 7.606666667 | 55.73 | 64.91666667 |
| **D.long019626** | 557 | Pfam | PF00232 | Glycosyl hydrolase family 1 | 6.49 | 8.703333333 | 10.32666667 | 9.886666667 |
| **D.long021797** | 120 | Pfam | PF00232 | Glycosyl hydrolase family 1 | 0 | 0 | 0 | 0 |
| **D.long021814** | 502 | Pfam | PF00232 | Glycosyl hydrolase family 1 | 0 | 0 | 0 | 0 |
| **D.long021816** | 511 | Pfam | PF00232 | Glycosyl hydrolase family 1 | 0.023333333 | 0 | 0 | 0 |
| **D.long021818** | 514 | Pfam | PF00232 | Glycosyl hydrolase family 1 | 0.813333333 | 0.563333333 | 0.046666667 | 12.5 |
| **D.long025080** | 514 | Pfam | PF00232 | Glycosyl hydrolase family 1 | 0.383333333 | 0 | 3.606666667 | 0 |
| **D.long025081** | 828 | Pfam | PF00232 | Glycosyl hydrolase family 1 | 11.63666667 | 0.59 | 30.6 | 2.04 |
| **D.long025824** | 315 | Pfam | PF00232 | Glycosyl hydrolase family 1 | 0 | 0.076666667 | 0 | 0 |
| **D.long025826** | 524 | Pfam | PF00232 | Glycosyl hydrolase family 1 | 1.866666667 | 0.036666667 | 0.466666667 | 0.453333333 |
| **D.long026156** | 524 | Pfam | PF00232 | Glycosyl hydrolase family 1 | 1.903333333 | 0 | 0.243333333 | 0.673333333 |
| **D.long026162** | 222 | Pfam | PF00232 | Glycosyl hydrolase family 1 | 0 | 0 | 0 | 0 |
| **D.long028671** | 618 | Pfam | PF00232 | Glycosyl hydrolase family 1 | 6.723333333 | 12.68 | 9.346666667 | 5.65 |
| **D.long028672** | 525 | Pfam | PF00232 | Glycosyl hydrolase family 1 | 0.576666667 | 49.13 | 7.92 | 3.413333333 |
| **D.long028674** | 535 | Pfam | PF00232 | Glycosyl hydrolase family 1 | 4.813333333 | 60.19 | 5.996666667 | 25.4 |
| **D.long028675** | 510 | Pfam | PF00232 | Glycosyl hydrolase family 1 | 0 | 5.316666667 | 1.533333333 | 14.73333333 |
| **D.long035562** | 161 | Pfam | PF00232 | Glycosyl hydrolase family 1 | 0.256666667 | 0 | 0 | 0.1 |
| **D.long035566** | 520 | Pfam | PF00232 | Glycosyl hydrolase family 1 | 6 | 40.46 | 5.683333333 | 14.08 |
| **D.long037807** | 500 | Pfam | PF00232 | Glycosyl hydrolase family 1 | 0.11 | 0.036666667 | 0.056666667 | 0.053333333 |
| **D.long037808** | 500 | Pfam | PF00232 | Glycosyl hydrolase family 1 | 0.773333333 | 0.043333333 | 0.766666667 | 0.406666667 |
| **D.long037809** | 494 | Pfam | PF00232 | Glycosyl hydrolase family 1 | 10.44 | 0 | 0.433333333 | 0.023333333 |
| **D.long001393** | 513 | PRINTS | PR00131 | Glycosyl hydrolase family 1 signature | 0.116666667 | 0 | 0 | 1.933333333 |
| **D.long009526** | 521 | PRINTS | PR00131 | Glycosyl hydrolase family 1 signature | 1.833333333 | 47.78666667 | 22.01333333 | 25.31333333 |
| **D.long010988** | 515 | PRINTS | PR00131 | Glycosyl hydrolase family 1 signature | 16.18666667 | 0 | 3.546666667 | 11.00333333 |
| **D.long011003** | 513 | PRINTS | PR00131 | Glycosyl hydrolase family 1 signature | 13.69666667 | 0.016666667 | 0.366666667 | 2.113333333 |
| **D.long019625** | 527 | PRINTS | PR00131 | Glycosyl hydrolase family 1 signature | 8.603333333 | 1.02 | 5.043333333 | 7.313333333 |
| **D.long032044** | 515 | PRINTS | PR00131 | Glycosyl hydrolase family 1 signature | 28.65333333 | 0 | 0.086666667 | 0.996666667 |
| **D.long004123** | 238 | Pfam | PF01596 | O-methyltransferase | 38.20333333 | 11.58333333 | 21.68333333 | 29.74666667 |
| **D.long004124** | 238 | Pfam | PF01596 | O-methyltransferase | 43.85 | 18.47666667 | 28.08 | 34.45 |
| **D.long004126** | 238 | Pfam | PF01596 | O-methyltransferase | 138.8866667 | 11.40333333 | 79.62666667 | 28.78666667 |
| **D.long000403** | 320 | Pfam | PF00141 | Peroxidase | 76.15666667 | 0.073333333 | 0.24 | 0.036666667 |
| **D.long000404** | 335 | Pfam | PF00141 | Peroxidase | 0 | 0 | 0 | 0 |
| **D.long000406** | 373 | Pfam | PF00141 | Peroxidase | 0 | 0 | 0 | 0 |
| **D.long004529** | 322 | Pfam | PF00141 | Peroxidase | 32.2 | 18.20666667 | 25.33666667 | 45.27333333 |
| **D.long006178** | 336 | Pfam | PF00141 | Peroxidase | 0 | 0 | 0 | 0 |
| **D.long006180** | 338 | Pfam | PF00141 | Peroxidase | 6.576666667 | 0.123333333 | 1.11 | 1.46 |
| **D.long006181** | 337 | Pfam | PF00141 | Peroxidase | 2.996666667 | 0 | 2.51 | 2.136666667 |
| **D.long006182** | 130 | Pfam | PF00141 | Peroxidase | 0 | 0 | 0 | 0 |
| **D.long006183** | 338 | Pfam | PF00141 | Peroxidase | 0.086666667 | 0 | 3.403333333 | 0.576666667 |
| **D.long006184** | 338 | Pfam | PF00141 | Peroxidase | 0.086666667 | 0 | 2.623333333 | 0.883333333 |
| **D.long006185** | 335 | Pfam | PF00141 | Peroxidase | 0 | 0.073333333 | 0.193333333 | 0.133333333 |
| **D.long006479** | 322 | Pfam | PF00141 | Peroxidase | 1.18 | 0.383333333 | 1.2 | 0 |
| **D.long012043** | 326 | Pfam | PF00141 | Peroxidase | 0 | 1.466666667 | 0.21 | 0.113333333 |
| **D.long012051** | 324 | Pfam | PF00141 | Peroxidase | 0.05 | 762.3133333 | 0.32 | 0 |
| **D.long012053** | 336 | Pfam | PF00141 | Peroxidase | 4.04 | 486.1566667 | 61.42 | 225.7366667 |
| **D.long012056** | 332 | Pfam | PF00141 | Peroxidase | 0 | 0.58 | 5.89 | 3.016666667 |
| **D.long012057** | 327 | Pfam | PF00141 | Peroxidase | 13.11 | 1.313333333 | 12.95333333 | 3.603333333 |
| **D.long012540** | 281 | Pfam | PF00141 | Peroxidase | 0 | 2.24 | 0.26 | 0 |
| **D.long014332** | 422 | Pfam | PF00141 | Peroxidase | 0.03 | 0 | 0 | 1.793333333 |
| **D.long014333** | 330 | Pfam | PF00141 | Peroxidase | 0 | 0 | 0 | 0 |
| **D.long014334** | 322 | Pfam | PF00141 | Peroxidase | 0 | 0 | 0 | 0 |
| **D.long024588** | 366 | Pfam | PF00141 | Peroxidase | 0 | 0 | 0 | 0 |
| **D.long025054** | 320 | Pfam | PF00141 | Peroxidase | 25.18666667 | 0 | 1.766666667 | 0.88 |
| **D.long025112** | 318 | Pfam | PF00141 | Peroxidase | 0.05 | 0 | 0 | 0 |
| **D.long029149** | 330 | Pfam | PF00141 | Peroxidase | 0 | 0 | 0 | 0 |
| **D.long029152** | 329 | Pfam | PF00141 | Peroxidase | 0.473333333 | 0 | 0 | 0.323333333 |
| **D.long029153** | 329 | Pfam | PF00141 | Peroxidase | 0.793333333 | 8.89 | 3.423333333 | 3.433333333 |
| **D.long032015** | 275 | Pfam | PF00141 | Peroxidase | 6.296666667 | 1.806666667 | 12.55 | 20.28333333 |
| **D.long032016** | 331 | Pfam | PF00141 | Peroxidase | 1.036666667 | 2.306666667 | 0.236666667 | 3.883333333 |
| **D.long034018** | 338 | Pfam | PF00141 | Peroxidase | 11.76 | 1.583333333 | 1.253333333 | 2.363333333 |
| **D.long035259** | 337 | Pfam | PF00141 | Peroxidase | 0.036666667 | 0 | 0 | 0 |
| **D.long035294** | 339 | Pfam | PF00141 | Peroxidase | 0 | 0 | 0.03 | 0.103333333 |
| **D.long037726** | 353 | Pfam | PF00141 | Peroxidase | 35.64333333 | 62.85666667 | 46.55 | 55.29333333 |
| **D.long037727** | 353 | Pfam | PF00141 | Peroxidase | 63.76666667 | 188.9866667 | 68.08666667 | 135.3533333 |
| **D.long037728** | 314 | Pfam | PF00141 | Peroxidase | 0 | 0 | 0 | 0.193333333 |
| **D.long037729** | 320 | Pfam | PF00141 | Peroxidase | 0.146666667 | 0.11 | 0 | 1.503333333 |
| **D.long037730** | 322 | Pfam | PF00141 | Peroxidase | 4.276666667 | 2.556666667 | 0.643333333 | 1.203333333 |
| **D.long037732** | 320 | Pfam | PF00141 | Peroxidase | 0 | 0 | 0 | 0 |
| **D.long005200** | 709 | TIGRFAM | TIGR01226 | phe_am_lyase: phenylalanine ammonia-lyase | 269.7766667 | 403.8066667 | 291.8966667 | 80.39666667 |
| **D.long007573** | 706 | TIGRFAM | TIGR01226 | phe_am_lyase: phenylalanine ammonia-lyase | 44.87 | 88.78333333 | 40.91333333 | 22.40666667 |
| **D.long007574** | 702 | TIGRFAM | TIGR01226 | phe_am_lyase: phenylalanine ammonia-lyase | 24.70666667 | 1.01 | 12.5 | 6.973333333 |
| **D.long007575** | 706 | TIGRFAM | TIGR01226 | phe_am_lyase: phenylalanine ammonia-lyase | 1.416666667 | 0 | 0.036666667 | 0.016666667 |
| **D.long034387** | 725 | TIGRFAM | TIGR01226 | phe_am_lyase: phenylalanine ammonia-lyase | 135.7666667 | 139.9366667 | 383.6133333 | 293.9133333 |

Table S7. List of different expressed IPR enriched gene families.

| #IPR id | IPR description |
|---|---|
| IPR002213 | UDP-glucuronosyl/UDP-glucosyltransferase |
| IPR002401 | Cytochrome P450, E-class, group I |
| IPR000719 | Protein kinase domain |
| IPR036396 | Cytochrome P450 superfamily |
| IPR001128 | Cytochrome P450 |
| IPR036852 | Peptidase S8/S53 domain superfamily |
| IPR000209 | Peptidase S8/S53 domain |
| IPR002160 | Proteinase inhibitor I3, Kunitz legume |
| IPR000858 | S-locus glycoprotein domain |
| IPR021720 | Malectin domain |
| IPR010259 | Peptidase S8 propeptide/proteinase inhibitor I9 |
| IPR000726 | Glycoside hydrolase, family 19, catalytic |
| IPR020843 | Polyketide synthase, enoylreductase domain |
| IPR025287 | Wall-associated receptor kinase, galacturonan-binding domain |
| IPR005123 | Oxoglutarate/iron-dependent dioxygenase |

Table S8. List of UGTs genes ID and their expression level.

| #transcript_id | gene_id | leaf | root | stem | fruit |
|---|---|---|---|---|---|
| **D.long000750.01** | D.long000750 | 14.56 | 0.18 | 17.91667 | 19.68 |
| **D.long000751.01** | D.long000751 | 3.97 | 3.035 | 4.113333 | 13.285 |
| **D.long000752.01** | D.long000752 | 0.82 | 33.32 | 7.256667 | 2.27 |
| **D.long000753.01** | D.long000753 | 24.74667 | 79.92 | 29.59667 | 33.4 |
| **D.long000754.01** | D.long000754 | 0.066667 | 30.06 | 0.49 | 0.385 |
| **D.long001253.01** | D.long001253 | 12.57 | 0.04 | 0.803333 | 0.03 |
| **D.long001256.01** | D.long001256 | 14.22333 | 7.055 | 1.383333 | 3.585 |
| **D.long002736.01** | D.long002736 | 5.796667 | 4.31 | 7.143333 | 12.69 |
| **D.long004697.01** | D.long004697 | 0.023333 | 0.04 | 0.506667 | 0.395 |
| **D.long005887.01** | D.long005887 | 13.21 | 18.305 | 21.78333 | 35.795 |
| **D.long005889.01** | D.long005889 | 7.253333 | 20.47 | 11.41667 | 20.53 |
| **D.long005890.01** | D.long005890 | 0.623333 | 14.395 | 3.55 | 5.945 |
| **D.long005892.01** | D.long005892 | 369.6867 | 313.02 | 183.1533 | 182.915 |
| **D.long007001.01** | D.long007001 | 0.26 | 3.44 | 0.773333 | 0 |
| **D.long007002.01** | D.long007002 | 15.17333 | 0.23 | 17.84 | 8.14 |
| **D.long007337.01** | D.long007337 | 0 | 0 | 0 | 0.12 |
| **D.long007914.01** | D.long007914 | 0.63 | 0 | 0.02 | 13.995 |
| **D.long007915.01** | D.long007915 | 0 | 0.04 | 0 | 1.285 |
| **D.long007934.01** | D.long007934 | 15.66 | 51.695 | 48.11667 | 37.99 |
| **D.long007936.01** | D.long007936 | 0.346667 | 4.42 | 5.473333 | 2.555 |
| **D.long007938.01** | D.long007938 | 5.53 | 3.545 | 0.753333 | 0.55 |
| **D.long007941.01** | D.long007941 | 0.303333 | 0.545 | 1.233333 | 1.38 |
| **D.long007942.01** | D.long007942 | 3.093333 | 0.855 | 13.17667 | 9.065 |
| **D.long007958.01** | D.long007958 | 0.226667 | 0 | 0 | 0.08 |
| **D.long007959.01** | D.long007959 | 0.066667 | 0.36 | 0.02 | 0 |
| **D.long008122.01** | D.long008122 | 0.046667 | 24.275 | 2.89 | 3.9 |
| **D.long008123.01** | D.long008123 | 0.02 | 5.495 | 1.283333 | 0.835 |
| **D.long008126.01** | D.long008126 | 0 | 0.95 | 0 | 0 |
| **D.long008130.01** | D.long008130 | 1.646667 | 0.505 | 0.46 | 1.635 |
| **D.long008131.01** | D.long008131 | 12.77 | 10.26 | 6.163333 | 13.235 |
| **D.long008459.01** | D.long008459 | 2.873333 | 23.325 | 5.913333 | 1.62 |
| **D.long009885.01** | D.long009885 | 0 | 0 | 0.053333 | 0 |
| **D.long011239.01** | D.long011239 | 56.57667 | 0.885 | 97.63 | 49.335 |
| **D.long013435.01** | D.long013435 | 0.316667 | 46.305 | 3.84 | 5.25 |
| **D.long015203.01** | D.long015203 | 0.243333 | 0 | 0 | 0.12 |
| **D.long015746.01** | D.long015746 | 8.88 | 7.09 | 1.14 | 2.2 |
| **D.long016471.01** | D.long016471 | 7.073333 | 8.7 | 10.35667 | 14.375 |
| **D.long016946.01** | D.long016946 | 0.406667 | 0.1 | 0.263333 | 0.085 |
| **D.long017794.01** | D.long017794 | 0.643333 | 0.93 | 0.303333 | 0.215 |
| **D.long019494.01** | D.long019494 | 3.79 | 24.95 | 68.02333 | 40.25 |
| **D.long019495.01** | D.long019495 | 11.6 | 0.22 | 3.193333 | 3.165 |
| **D.long019496.01** | D.long019496 | 0 | 27.97 | 0.063333 | 0.055 |
| **D.long019506.01** | D.long019506 | 0 | 2.405 | 1.1 | 0.25 |
| **D.long019508.01** | D.long019508 | 0.146667 | 6.765 | 1.443333 | 0.93 |
| **D.long019513.01** | D.long019513 | 0.11 | 16.505 | 0 | 0.135 |
| **D.long020005.01** | D.long020005 | 0 | 0.05 | 0 | 0 |
| **D.long020006.01** | D.long020006 | 0 | 4.515 | 0.883333 | 0.265 |
| **D.long020009.01** | D.long020009 | 0 | 1.45 | 0.823333 | 0.725 |
| **D.long022372.01** | D.long022372 | 0.016667 | 0 | 0.443333 | 0.85 |
| **D.long023017.01** | D.long023017 | 0.18 | 0.04 | 0.083333 | 0.025 |

| | | | | | |
|---|---|---|---|---|---|
| **D.long023291.01** | D.long023291 | 1.1 | 16.33 | 5.62 | 6.15 |
| **D.long023303.01** | D.long023303 | 31.83 | 30.09 | 58.61 | 70.03 |
| **D.long023305.01** | D.long023305 | 0.07 | 0 | 0.443333 | 2.63 |
| **D.long023307.01** | D.long023307 | 0 | 0 | 0.02 | 0.285 |
| **D.long023309.01** | D.long023309 | 0.523333 | 21.8 | 2.81 | 63.75 |
| **D.long024681.01** | D.long024681 | 3.493333 | 0.07 | 0.17 | 0 |
| **D.long026389.01** | D.long026389 | 1.073333 | 12.585 | 5.703333 | 4.31 |
| **D.long027024.01** | D.long027024 | 0.436667 | 4.46 | 5.92 | 6.515 |
| **D.long027795.01** | D.long027795 | 0 | 0 | 0.226667 | 0 |
| **D.long027797.01** | D.long027797 | 0.763333 | 19.455 | 1.323333 | 1.8 |
| **D.long027798.01** | D.long027798 | 22.66667 | 8.01 | 16.66 | 12.64 |
| **D.long030886.01** | D.long030886 | 23.56333 | 79.455 | 78.65 | 60.97 |
| **D.long030887.01** | D.long030887 | 5.773333 | 30.575 | 58.18667 | 34.54 |
| **D.long033649.01** | D.long033649 | 0.553333 | 3.045 | 1.306667 | 8.08 |
| **D.long033685.01** | D.long033685 | 0.033333 | 5.245 | 0 | 3.73 |
| **D.long033689.01** | D.long033689 | 0.793333 | 10.27 | 1.626667 | 0.175 |
| **D.long033693.01** | D.long033693 | 0.296667 | 2.835 | 0.156667 | 4.745 |
| **D.long033697.01** | D.long033697 | 0.14 | 1.975 | 0.193333 | 0.86 |
| **D.long033704.01** | D.long033704 | 6.91 | 25.825 | 10.51667 | 0.4 |
| **D.long033705.01** | D.long033705 | 5.44 | 1.66 | 21.44333 | 33.305 |
| **D.long033707.01** | D.long033707 | 1.573333 | 3.825 | 14.85 | 47.005 |
| **D.long033718.01** | D.long033718 | 0.583333 | 1.095 | 7.45 | 8.475 |
| **D.long033719.01** | D.long033719 | 146.17 | 0 | 11.24333 | 1.48 |
| **D.long033721.01** | D.long033721 | 1.393333 | 0.115 | 0 | 0.025 |
| **D.long034281.01** | D.long034281 | 0.14 | 1.835 | 0.603333 | 0.345 |
| **D.long034282.01** | D.long034282 | 0 | 0.035 | 0 | 0.075 |
| **D.long034283.01** | D.long034283 | 0.29 | 31 | 3.61 | 13.985 |
| **D.long034284.01** | D.long034284 | 9.436667 | 12.545 | 9.266667 | 13.795 |
| **D.long034285.01** | D.long034285 | 0 | 0 | 0.056667 | 0.355 |
| **D.long034286.01** | D.long034286 | 3.4 | 0 | 3.013333 | 3.575 |
| **D.long034287.01** | D.long034287 | 0.06 | 0 | 0.053333 | 0 |
| **D.long034289.01** | D.long034289 | 1.396667 | 0.855 | 13.27 | 10.35 |
| **D.long034290.01** | D.long034290 | 0.64 | 4.975 | 6.203333 | 2.355 |
| **D.long034371.01** | D.long034371 | 0.02 | 1.06 | 0 | 0.335 |
| **D.long036345.01** | D.long036345 | 0.146667 | 0.19 | 0.07 | 12.725 |
| **D.long036346.01** | D.long036346 | 23.62333 | 65.915 | 51.52333 | 42.67 |
| **D.long036348.01** | D.long036348 | 0.023333 | 0.695 | 0.06 | 29.235 |
| **D.long036350.01** | D.long036350 | 0.64 | 0.39 | 0.15 | 4.395 |
| **D.long036353.01** | D.long036353 | 0.316667 | 6.125 | 0.69 | 20.27 |
| **D.long036354.01** | D.long036354 | 0.066667 | 0.495 | 0.233333 | 7.195 |
| **D.long036357.01** | D.long036357 | 1.173333 | 1.8 | 1.616667 | 2.845 |
| **D.long036906.01** | D.long036906 | 0 | 0.96 | 0 | 4.9 |
| **D.long036908.01** | D.long036908 | 0.423333 | 1.46 | 0.596667 | 14.105 |
| **D.long036910.01** | D.long036910 | 0.653333 | 7 | 3.053333 | 8.095 |
| **D.long037404.01** | D.long037404 | 0 | 0 | 0 | 0.135 |
| **D.long037439.01** | D.long037439 | 1.433333 | 2.83 | 2.046667 | 0.175 |

**Table S9. List of genome resequencing samples and their locations.**

| Sequencing-ID | First letter of cultivar and its location | Location (Province/Country) | City | Artificial Breeding |
|---|---|---|---|---|
| D01 | CL-GD | GuangDong | GaoZhou | No |
| D02 | SMM-GD | GuangDong | GaoZhou | No |
| D03 | BHM-GD | GuangDong | GaoZhou | No |
| D04 | LSM-GD | GuangDong | GaoZhou | No |
| D05 | TBM-GD | GuangDong | GaoZhou | No |
| D06 | SLM-GD | GuangDong | GaoZhou | No |
| D07 | HH-GD | GuangDong | GaoZhou | No |
| D08 | GSEH-GD | GuangDong | JieYang | No |
| D09 | WGTGY-GD | GuangDong | GaoZhou | No |
| D10 | JRY-GD | GuangDong | GaoZhou | No |
| D11 | HLGY-GD | GuangDong | GaoZhou | No |
| D12 | JSY-GD | GuangDong | GaoZhou | No |
| D13 | SLR-GD | GuangDong | GaoZhou | No |
| D14 | HDGY-GD | GuangDong | GaoZhou | No |
| D15 | YC-GD | GuangDong | GuangZhou | Yes |
| D16 | GZ-GD | GuangDong | GuangZhou | Yes |
| D17 | ZBL-GX | GuangXi | GuiPing | No |
| D18 | SSCR-GD | GuangDong | GuangZhou | Yes |
| D19 | CHDGY-GD | GuangDong | CongHua | No |
| D20 | XJWY-GD | GuangDong | GuangZhou | Yes |
| D21 | LZZ-GD | GuangDong | JieYang | No |
| D22 | SSCS-GD | GuangDong | GuangZhou | Yes |
| D23 | XJYH-GD | GuangDong | GuangZhou | Yes |
| D24 | SX-GD | GuangDong | NanHai | No |
| D25 | SY-GD | GuangDong | ZhongShan | No |
| D26 | DWY-GX | GuangXi | YuLin | No |
| D27 | JY-GD | GuangDong | GuangZhou | Yes |
| D28 | KHL-Aus | Australia | unknown | No |
| D29 | YD-Thai | Thailand | unknown | No |
| D30 | PT-Thai | Thailand | unknown | No |
| D31 | XY-GD | GuangDong | GuangZhou | Yes |
| D32 | TXZQ-GX | GuangXi | WuZhou | No |
| D33 | YZDWY-GX | GuangXi | YuLin | No |
| D34 | GZA1-GD | GuangDong | GuangZhou | Yes |
| D35 | GZ3-GD | GuangDong | GuangZhou | Yes |
| D36 | DWYH-GD | GuangDong | DongGuan | No |
| D37 | DF-GD | GuangDong | DongGuan | No |
| D38 | LY-GD | GuangDong | GuangZhou | Yes |
| D39 | TGSJ-Thai | Thailand | unknown | No |
| D40 | YTB-FJ | FuJian | PuTian | No |
| D41 | SZP-Thai | Thailand | unknown | No |
| D42 | JL-FJ | FuJian | PuTian | No |
| D43 | HXB-FJ | FuJian | PuTian | No |
| D44 | QKBY-FJ | FuJian | ChangLe | No |
| D45 | SZ-FJ | FuJian | XiaMen | No |
| D46 | WCL-FJ | FuJian | ChangLe | No |
| D47 | PMA-FJ | FuJian | PuTian | No |
| D48 | GX-GX | GuangXi | NanNing | Yes |
| D49 | SG-SC | SiChuan | LuZhou | No |
| D50 | JYW-FJ | FuJian | PuTian | No |
| D51 | LY-FJ | FuJian | PuTian | No |
| D52 | LDB-FJ | FuJian | PuTian | No |
| D54 | GLZ-GX | GuangXi | NanNing | No |
| D55 | LF-SC | SiChuan | LuZhou | No |
| D56 | CHZ-GD | GuangDong | JieYang | No |
| D57 | CPZ-GD | GuangDong | RaoPing | No |
| D58 | CK-FJ | FuJian | XiaMen | No |
| D59 | GMB-FJ | FuJian | PuTian | No |
| D60 | SNYH-FJ | FuJian | PuTian | No |
| D61 | HBP-FJ | FuJian | TongAn | No |
| D62 | HKZ-FJ | FuJian | unknown | No |
| D63 | HY-GD | GuangDong | Taishan | No |
| D64 | CSB-FJ | FuJian | PuTian | No |
| D65 | QZB-FJ | FuJian | PuTian | No |
| D66 | WLL-FJ | FuJian | XianYou | No |
| D67 | JDB-FJ | FuJian | PuTian | No |
| D68 | ZTB-FJ | FuJian | QuanZhou | No |
| D69 | LQB-FJ | FuJian | PuTian | No |
| D70 | FLD | GuangDong | ChaoZhou | No |
| D71 | BYZ-FJ | FuJian | TongAn | No |
| D72 | YY106-FJ | FuJian | PuTian | No |
| D73 | QKJ-FJ | FuJian | unknown | No |
| D74 | SFB-FJ | FuJian | PuTian | No |
| D75 | GHW | GuangDong | GuangZhou | Yes |
| D76 | FY-FJ | FuJian | FuZhou | No |
| D77 | JH-GD | GuangDong | GuangZhou | Yes |
| D78 | LZ-SC | SiChuan | LuZhou | No |
| D79 | GZA2-GD | GuangDong | GuangZhou | Yes |
| D80 | YNSJ-Viet | Vietnam | unknown | No |
| D81 | FLS-FJ | FuJian | XiaMen | No |
| D82 | FLHK-SC | SiChuan | ChongQing | No |
| D83 | FDWM-GD | GuangDong | ChaoZhou | Yes |
| D84 | LY-HN | HaiNan | unknown | No |
| D85 | GZ2-GD | GuangDong | GuangZhou | Yes |
| D86 | DB-FJ | FuJian | QuanZhou | No |
| D87 | GMYH-GX | GuangXi | NanNing | Yes |
| D88 | HXZ-FJ | FuJian | ChangLe | No |

Table S10. GWAS results of significant SNPs associated with six traits and their annotation.

| Trait | Chromosome | Gene id | Promoter | Intron | CDS | Synonymous | Nonsynonymous |
|---|---|---|---|---|---|---|---|
| Pericarp Thickness | chr1 | D.long001288.01 | 7 | 0 | 0 | 0 | 0 |
| | | D.long001288.02 | 0 | 46 | 6 | 3 | 3 |
| | | D.long001289.01 | 13 | 141 | 20 | 14 | 6 |
| | | D.long001290.01 | 12 | 18 | 4 | 2 | 2 |
| | | D.long001291.01 | 1 | 23 | 0 | 0 | 0 |
| | | D.long001292.01 | 2 | 106 | 26 | 10 | 16 |
| | | D.long001293.01 | 27 | 0 | 266 | 58 | 208 |
| | | D.long001294.01 | 70 | 13 | 109 | 34 | 75 |
| | | D.long001295.01 | 10 | 0 | 69 | 22 | 47 |
| | | D.long001296.01 | 1 | 108 | 273 | 273 | 0 |
| | | D.long001297.01 | 11 | 0 | 41 | 8 | 33 |
| | | D.long001298.01 | 33 | 10 | 55 | 16 | 39 |
| | | D.long001299.01 | 10 | 240 | 34 | 11 | 23 |
| | | D.long001300.01 | 2 | 11 | 35 | 8 | 27 |
| | | D.long001301.01 | 16 | 75 | 6 | 2 | 4 |
| | | D.long001302.01 | 4 | 54 | 8 | 2 | 6 |
| | | D.long001303.01 | 3 | 68 | 7 | 2 | 5 |
| | | D.long001304.01 | 5 | 0 | 14 | 6 | 8 |
| | | D.long001305.01 | 23 | 0 | 17 | 8 | 9 |
| | | D.long001306.01 | 20 | 110 | 5 | 1 | 4 |
| | | D.long001307.01 | 30 | 9 | 31 | 11 | 20 |
| | | D.long001308.01 | 1 | 19 | 12 | 3 | 9 |
| | | D.long001309.01 | 5 | 2 | 1 | 1 | 0 |
| | | D.long001310.01 | 25 | 15 | 5 | 2 | 3 |
| | | D.long001311.01 | 14 | 60 | 15 | 8 | 7 |
| | chr6 | D.long025467.01 | 39 | 2 | 23 | 10 | 13 |
| | | D.long025468.01 | 11 | 0 | 27 | 11 | 16 |
| | | D.long025469.01 | 38 | 32 | 85 | 36 | 49 |
| | | D.long025470.01 | 4 | 18 | 5 | 2 | 3 |
| | | D.long025471.01 | 0 | 144 | 44 | 14 | 30 |
| | | D.long025472.01 | 19 | 0 | 13 | 4 | 9 |
| | | D.long025473.01 | 1 | 30 | 51 | 15 | 36 |
| | | D.long025474.01 | 60 | 11 | 68 | 14 | 54 |
| | | D.long025475.01 | 18 | 56 | 11 | 10 | 1 |
| | | D.long025476.01 | 5 | 4 | 23 | 6 | 17 |
| | | D.long025477.01 | 3 | 66 | 28 | 11 | 17 |

| Trait | Chromosome | Gene id | Promoter | Intron | CDS | Synonymous | Nonsynonymous |
|---|---|---|---|---|---|---|---|
| **Pulp Thickness** | chr5 | D.long015104.01 | 17 | 4 | 14 | 5 | 9 |
| | | D.long015105.01 | 15 | 1 | 84 | 29 | 55 |
| | | D.long015106.01 | 40 | 6 | 42 | 30 | 12 |
| | | D.long015107.01 | 0 | 0 | 56 | 56 | 0 |
| | | D.long015108.01 | 31 | 11 | 36 | 14 | 22 |
| | | D.long015109.01 | 5 | 29 | 42 | 10 | 32 |
| | | D.long015110.01 | 10 | 70 | 44 | 9 | 35 |
| | | D.long015111.01 | 6 | 5 | 32 | 8 | 24 |
| | | D.long015112.01 | 28 | 13 | 30 | 8 | 22 |
| | | D.long015113.01 | 29 | 0 | 42 | 10 | 32 |
| | | D.long015114.01 | 0 | 0 | 29 | 18 | 11 |
| | | D.long015115.01 | 27 | 0 | 17 | 2 | 15 |
| | | D.long015116.01 | 5 | 0 | 4 | 1 | 3 |
| | | D.long015117.01 | 24 | 7 | 20 | 5 | 15 |
| | | D.long015118.01 | 20 | 7 | 17 | 10 | 7 |

| Trait | Chromosome | Gene id | Promoter | Intron | CDS | Synonymous | Nonsynonymous |
|---|---|---|---|---|---|---|---|
| **Fruit' Horizontal Diameter** | chr10 | D.long035939.01 | 3 | 18 | 6 | 5 | 1 |
| | | D.long035940.01 | 1 | 32 | 7 | 7 | 0 |
| | | D.long035941.01 | 4 | 0 | 18 | 4 | 14 |
| | | D.long035942.01 | 14 | 8 | 21 | 6 | 15 |
| | | D.long035943.01 | 17 | 57 | 54 | 14 | 40 |
| | | D.long035944.01 | 1 | 14 | 0 | 0 | 0 |
| | | D.long035945.01 | 6 | 1 | 5 | 0 | 5 |
| | | D.long035946.01 | 14 | 0 | 5 | 3 | 2 |
| | | D.long035947.01 | 1 | 81 | 7 | 3 | 4 |
| | | D.long035948.01 | 14 | 0 | 7 | 3 | 4 |
| | | D.long035949.01 | 5 | 8 | 12 | 3 | 9 |
| | | D.long035950.02 | 14 | 1 | 7 | 3 | 4 |
| | chr11 | D.long027632.01 | 15 | 7 | 6 | 4 | 2 |
| | | D.long027633.01 | 2 | 5 | 6 | 2 | 4 |
| | | D.long027634.01 | 2 | 0 | 1 | 0 | 1 |
| | | D.long027635.01 | 5 | 1 | 3 | 2 | 1 |
| | | D.long027636.01 | 9 | 1 | 6 | 1 | 5 |
| | | D.long027637.01 | 0 | 0 | 1 | 0 | 1 |
| | | D.long027638.01 | 0 | 0 | 1 | 0 | 1 |
| | | D.long027639.01 | 5 | 3 | 0 | 0 | 0 |
| | | D.long027640.01 | 0 | 2 | 2 | 0 | 2 |
| | | D.long027641.01 | 3 | 3 | 0 | 0 | 0 |
| | | D.long027642.01 | 0 | 0 | 1 | 1 | 0 |
| | | D.long027643.01 | 9 | 2 | 12 | 5 | 7 |
| | | D.long027644.01 | 44 | 2 | 38 | 7 | 31 |
| | | D.long027645.01 | 0 | 0 | 2 | 0 | 2 |
| | | D.long027646.01 | 65 | 89 | 97 | 97 | 0 |
| | | D.long027647.01 | 11 | 39 | 16 | 11 | 5 |
| | | D.long027648.01 | 0 | 1 | 2 | 1 | 1 |
| | | D.long027649.01 | 12 | 0 | 2 | 2 | 0 |
| | | D.long027650.01 | 3 | 0 | 6 | 4 | 2 |
| | | D.long027651.01 | 3 | 9 | 10 | 4 | 6 |
| | | D.long027652.01 | 4 | 49 | 9 | 2 | 7 |
| | | D.long027653.01 | 14 | 0 | 14 | 5 | 9 |
| | | D.long027654.01 | 11 | 0 | 14 | 4 | 10 |
| | | D.long027655.01 | 7 | 0 | 4 | 0 | 4 |
| | | D.long027656.01 | 4 | 2 | 11 | 5 | 6 |
| | | D.long027657.01 | 5 | 0 | 7 | 1 | 6 |

| Trait | Chromosome | Gene id | Promoter | Intron | CDS | Synonymous | Nonsynonymous |
|---|---|---|---|---|---|---|---|
| **Edible Rate** | chr14 | D.long030308.01 | 0 | 4 | 8 | 5 | 3 |
| | | D.long030309.01 | 37 | 137 | 55 | 20 | 35 |
| | | D.long030310.01 | 3 | 0 | 7 | 4 | 3 |
| | | D.long030311.01 | 24 | 0 | 13 | 6 | 7 |
| | | D.long030312.01 | 12 | 3 | 26 | 9 | 17 |
| | | D.long030313.01 | 30 | 0 | 37 | 17 | 20 |
| | | D.long030314.01 | 44 | 24 | 20 | 15 | 5 |
| | | D.long030315.01 | 22 | 11 | 14 | 8 | 6 |
| | | D.long030316.01 | 37 | 63 | 30 | 16 | 14 |
| | | D.long030317.01 | 7 | 125 | 12 | 4 | 8 |
| | | D.long030318.01 | 0 | 0 | 11 | 3 | 8 |
| | | D.long030320.01 | 8 | 43 | 19 | 5 | 14 |
| | | D.long030321.01 | 6 | 16 | 6 | 6 | 0 |
| | | D.long030322.01 | 26 | 17 | 0 | 0 | 0 |
| | | D.long030323.01 | 6 | 0 | 13 | 3 | 10 |
| | | D.long030324.01 | 15 | 320 | 26 | 10 | 16 |
| | | D.long030325.01 | 21 | 23 | 6 | 5 | 1 |
| | | D.long030326.01 | 4 | 0 | 32 | 9 | 23 |
| | | D.long030327.01 | 0 | 27 | 12 | 2 | 10 |
| | | D.long030328.01 | 5 | 0 | 2 | 0 | 2 |
| | | D.long030329.01 | 20 | 110 | 12 | 2 | 10 |

| Trait | Chromosome | Gene id | Promoter | Intron | CDS | Synonymous | Nonsynonymous |
|---|---|---|---|---|---|---|---|
| **Seed Weight** | chr12 | D.long038987.01 | 6 | 1 | 3 | 2 | 1 |
| | | D.long038988.01 | 15 | 0 | 11 | 3 | 8 |
| | | D.long038989.01 | 11 | 67 | 28 | 5 | 23 |
| | | D.long038990.01 | 9 | 0 | 16 | 8 | 8 |
| | | D.long038991.01 | 0 | 34 | 11 | 3 | 8 |
| | | D.long038992.01 | 40 | 62 | 68 | 20 | 48 |
| | | D.long038993.01 | 2 | 0 | 6 | 2 | 4 |
| | | D.long038994.01 | 5 | 0 | 4 | 1 | 3 |
| | | D.long038995.01 | 14 | 0 | 22 | 5 | 17 |
| | | D.long038996.01 | 12 | 30 | 42 | 7 | 35 |
| | | D.long038997.01 | 4 | 0 | 9 | 1 | 8 |
| | | D.long038998.01 | 28 | 13 | 64 | 22 | 42 |
| | | D.long038999.01 | 31 | 72 | 60 | 22 | 38 |
| | | D.long039000.01 | 1 | 0 | 1 | 0 | 1 |
| | | D.long039001.01 | 15 | 5 | 21 | 5 | 16 |
| | | D.long039002.01 | 1 | 9 | 0 | 0 | 0 |
| | | D.long039003.01 | 42 | 0 | 59 | 14 | 45 |
| | | D.long039004.01 | 63 | 105 | 197 | 75 | 122 |
| | | D.long039005.01 | 1 | 0 | 0 | 0 | 0 |
| | chr3 | D.long019821.01 | 11 | 46 | 8 | 3 | 5 |
| | | D.long019822.01 | 5 | 5 | 4 | 2 | 2 |
| | | D.long019823.01 | 13 | 33 | 19 | 9 | 10 |
| | | D.long019824.01 | 10 | 98 | 16 | 9 | 7 |
| | | D.long019825.01 | 49 | 10 | 90 | 72 | 18 |
| | | D.long019826.01 | 9 | 50 | 62 | 18 | 44 |
| | | D.long019827.01 | 11 | 12 | 4 | 3 | 1 |
| | | D.long019828.01 | 10 | 0 | 16 | 10 | 6 |
| | | D.long019829.01 | 1 | 2 | 1 | 0 | 1 |
| | | D.long019830.01 | 1 | 0 | 1 | 0 | 1 |
| | | D.long019831.01 | 28 | 77 | 2 | 1 | 1 |
| | | D.long019832.01 | 11 | 0 | 18 | 18 | 0 |
| | | D.long019833.01 | 19 | 0 | 31 | 31 | 0 |
| | | D.long019834.01 | 28 | 67 | 11 | 1 | 10 |
| | | D.long019835.01 | 0 | 4 | 1 | 0 | 1 |
| | | D.long019836.01 | 1 | 12 | 2 | 2 | 0 |
| | | D.long019837.01 | 8 | 33 | 24 | 14 | 10 |
| | | D.long019838.01 | 8 | 0 | 19 | 7 | 12 |
| | | D.long019839.01 | 38 | 49 | 49 | 20 | 29 |
| | | D.long019840.01 | 63 | 9 | 32 | 15 | 17 |
| | | D.long019841.01 | 12 | 29 | 13 | 3 | 10 |
| | | D.long019842.01 | 10 | 23 | 12 | 12 | 0 |
| | | D.long019843.01 | 7 | 7 | 52 | 17 | 35 |
| | | D.long019844.01 | 9 | 5 | 25 | 9 | 16 |
| | | D.long019845.01 | 8 | 36 | 2 | 1 | 1 |
| | | D.long019846.01 | 53 | 138 | 107 | 42 | 65 |
| | | D.long019847.01 | 12 | 140 | 6 | 3 | 3 |
| | | D.long019848.01 | 8 | 8 | 9 | 4 | 5 |
| | | D.long019849.01 | 0 | 1 | 1 | 1 | 0 |
| | chr6 | D.long024180.01 | 25 | 426 | 21 | 9 | 12 |
| | | D.long024181.01 | 35 | 58 | 5 | 1 | 4 |
| | | D.long024182.01 | 30 | 75 | 6 | 5 | 1 |
| | | D.long024183.01 | 5 | 382 | 3 | 0 | 3 |
| | | D.long024184.01 | 8 | 16 | 5 | 3 | 2 |
| | | D.long024185.01 | 7 | 131 | 92 | 42 | 50 |
| | | D.long024186.01 | 21 | 0 | 24 | 11 | 13 |
| | | D.long024187.01 | 49 | 147 | 23 | 7 | 16 |
| | | D.long024188.01 | 11 | 150 | 62 | 15 | 47 |
| | | D.long024189.01 | 24 | 114 | 18 | 6 | 12 |
| | | D.long024190.01 | 25 | 0 | 12 | 8 | 4 |
| | | D.long024191.01 | 12 | 195 | 14 | 4 | 10 |
| | | D.long024192.01 | 7 | 57 | 22 | 14 | 8 |
| | | D.long024193.01 | 17 | 0 | 2 | 0 | 2 |
| | | D.long024194.01 | 5 | 0 | 5 | 2 | 3 |
| | | D.long024130.01 | 14 | 0 | 104 | 39 | 65 |
| | | D.long024131.01 | 9 | 129 | 52 | 12 | 40 |
| | | D.long024132.01 | 35 | 23 | 48 | 16 | 32 |
| | | D.long024133.01 | 16 | 4 | 13 | 2 | 11 |
| | | D.long024134.01 | 1 | 24 | 14 | 2 | 12 |
| | | D.long024135.01 | 0 | 34 | 21 | 5 | 16 |
| | | D.long024135.02 | 9 | 0 | 0 | 0 | 0 |
| | | D.long024136.01 | 9 | 483 | 178 | 49 | 129 |
| | | D.long024137.01 | 13 | 159 | 33 | 11 | 22 |
| | | D.long024138.01 | 13 | 52 | 4 | 3 | 1 |
| | | D.long024139.01 | 4 | 9 | 0 | 0 | 0 |
| | | D.long024140.01 | 0 | 0 | 2 | 0 | 2 |
| | | D.long024141.01 | 9 | 51 | 6 | 6 | 0 |
| | | D.long024142.01 | 28 | 7 | 9 | 6 | 3 |
| | | D.long024143.01 | 0 | 22 | 96 | 33 | 63 |
| | | D.long024143.02 | 26 | 0 | 0 | 0 | 0 |
| | | D.long024144.01 | 10 | 44 | 26 | 13 | 13 |
| | | D.long024145.01 | 1 | 25 | 8 | 5 | 3 |
| | | D.long024146.01 | 7 | 51 | 21 | 8 | 13 |
| | | D.long024147.01 | 11 | 55 | 6 | 1 | 5 |
| | | D.long024148.01 | 38 | 75 | 16 | 3 | 13 |
| | | D.long024149.01 | 18 | 164 | 20 | 12 | 8 |
| | | D.long024150.01 | 3 | 81 | 27 | 8 | 19 |
| | | D.long024151.01 | 7 | 62 | 25 | 7 | 18 |
| | | D.long024152.01 | 30 | 321 | 38 | 16 | 22 |
| | | D.long024153.01 | 10 | 4 | 19 | 11 | 8 |
| | | D.long024154.01 | 10 | 7 | 0 | 0 | 0 |
| | | D.long024155.01 | 46 | 34 | 18 | 7 | 11 |
| | | D.long024156.01 | 21 | 187 | 76 | 17 | 59 |
| | | D.long024157.01 | 27 | 3 | 4 | 0 | 4 |
| | | D.long024158.01 | 16 | 32 | 21 | 5 | 16 |

| Trait | Chromosome | Gene id | Promoter | Intron | CDS | Synonymous | Nonsynonymous |
|---|---|---|---|---|---|---|---|
| **Total Soluble Solid** | chr14 | D.long029570.01 | 3 | 0 | 4 | 2 | 2 |
| | | D.long029571.01 | 5 | 17 | 12 | 3 | 9 |
| | | D.long029572.01 | 5 | 4 | 7 | 3 | 4 |
| | | D.long029573.01 | 3 | 0 | 1 | 1 | 0 |
| | | D.long029574.01 | 4 | 1 | 23 | 9 | 14 |
| | | D.long029575.01 | 14 | 4 | 6 | 1 | 5 |
| | | D.long029576.01 | 22 | 29 | 11 | 8 | 3 |
| | | D.long029577.01 | 22 | 36 | 7 | 2 | 5 |
| | | D.long029578.01 | 2 | 20 | 4 | 1 | 3 |
| | | D.long029579.01 | 12 | 19 | 4 | 3 | 1 |
| | | D.long029580.01 | 6 | 0 | 1 | 0 | 1 |
| | | D.long029581.01 | 3 | 10 | 4 | 4 | 0 |
| | | D.long029582.01 | 6 | 15 | 4 | 0 | 4 |
| | | D.long029583.01 | 7 | 0 | 2 | 0 | 2 |
| | | D.long029584.01 | 7 | 15 | 1 | 1 | 0 |
| | | D.long029585.01 | 3 | 112 | 6 | 1 | 5 |
| | | D.long029586.01 | 6 | 3 | 3 | 0 | 3 |
| | | D.long029587.01 | 12 | 28 | 15 | 6 | 9 |
| | | D.long029588.01 | 5 | 10 | 1 | 1 | 0 |
| | | D.long029589.01 | 31 | 89 | 23 | 12 | 11 |
| | | D.long029590.01 | 6 | 66 | 13 | 3 | 10 |
| | | D.long029591.01 | 20 | 77 | 20 | 6 | 14 |
| | | D.long029592.01 | 8 | 176 | 27 | 20 | 7 |
| | | D.long029593.01 | 8 | 58 | 14 | 5 | 9 |
| | | D.long029594.01 | 2 | 0 | 1 | 0 | 1 |
| | | D.long029595.01 | 3 | 9 | 4 | 3 | 1 |
| | | D.long029596.01 | 5 | 62 | 45 | 44 | 1 |
| | | D.long029597.01 | 9 | 0 | 0 | 0 | 0 |
| | | D.long029597.02 | 0 | 5 | 1 | 0 | 1 |
| | | D.long029598.01 | 0 | 6 | 3 | 1 | 2 |
| | | D.long029599.01 | 2 | 19 | 5 | 2 | 3 |
| | | D.long029600.01 | 2 | 1 | 0 | 0 | 0 |
| | | D.long029601.01 | 7 | 16 | 7 | 4 | 3 |
| | | D.long029602.01 | 0 | 12 | 8 | 3 | 5 |
| | | D.long029603.01 | 1 | 16 | 3 | 2 | 1 |
| | | D.long029604.01 | 4 | 13 | 6 | 1 | 5 |
| | | D.long029605.01 | 3 | 3 | 2 | 2 | 0 |
| | | D.long029606.01 | 2 | 0 | 6 | 4 | 2 |
| | | D.long029607.01 | 0 | 2 | 1 | 0 | 1 |
| | | D.long029608.01 | 3 | 0 | 0 | 0 | 0 |
| | | D.long029609.01 | 1 | 9 | 0 | 0 | 0 |
| | | D.long029610.01 | 2 | 0 | 4 | 3 | 1 |
| | | D.long029611.01 | 4 | 3 | 4 | 3 | 1 |
| | | D.long029612.01 | 8 | 5 | 10 | 4 | 6 |
| | | D.long029613.01 | 4 | 34 | 2 | 2 | 0 |
| | | D.long029614.01 | 10 | 0 | 0 | 0 | 0 |
| | chr3 | D.long019020.01 | 36 | 184 | 18 | 5 | 13 |
| | | D.long019021.01 | 15 | 54 | 9 | 6 | 3 |
| | | D.long019022.01 | 15 | 125 | 24 | 13 | 11 |
| | | D.long019023.01 | 1 | 104 | 22 | 5 | 17 |
| | | D.long019024.01 | 12 | 2 | 10 | 2 | 8 |
| | | D.long019025.01 | 20 | 10 | 44 | 15 | 29 |
| | | D.long019026.01 | 4 | 0 | 24 | 10 | 14 |
| | | D.long019027.01 | 5 | 79 | 17 | 6 | 11 |
| | | D.long019028.01 | 9 | 62 | 15 | 7 | 8 |
| | | D.long019029.01 | 33 | 81 | 55 | 23 | 32 |
| | | D.long019030.01 | 32 | 159 | 17 | 6 | 11 |
| | | D.long019031.01 | 13 | 227 | 29 | 12 | 17 |
| | | D.long019032.01 | 0 | 8 | 8 | 4 | 4 |
| | | D.long019033.01 | 17 | 24 | 43 | 12 | 31 |
| | | D.long019034.01 | 23 | 52 | 33 | 10 | 23 |
| | | D.long019035.01 | 2 | 25 | 105 | 45 | 60 |
| | | D.long019036.01 | 19 | 93 | 50 | 11 | 39 |
| | | D.long019801.01 | 10 | 26 | 4 | 2 | 2 |
| | | D.long019802.01 | 7 | 0 | 9 | 3 | 6 |
| | | D.long019803.01 | 6 | 49 | 23 | 17 | 6 |
| | | D.long019804.01 | 3 | 1 | 30 | 12 | 18 |
| | | D.long019805.01 | 6 | 0 | 6 | 3 | 3 |
| | | D.long019806.01 | 8 | 0 | 13 | 4 | 9 |
| | | D.long019807.01 | 0 | 0 | 21 | 10 | 11 |
| | | D.long019808.01 | 7 | 0 | 6 | 0 | 6 |
| | | D.long019810.01 | 4 | 12 | 3 | 1 | 2 |
| | | D.long019811.01 | 2 | 9 | 13 | 2 | 11 |
| | | D.long019812.01 | 14 | 14 | 29 | 13 | 16 |
| | | D.long019813.01 | 31 | 4 | 49 | 16 | 33 |
| | | D.long019814.01 | 3 | 3 | 19 | 6 | 13 |
| | | D.long019816.01 | | 11 | 18 | 5 | 13 |
| | | D.long019817.01 | 4 | 38 | 20 | 2 | 18 |
| | | D.long019818.01 | 8 | 111 | 36 | 26 | 10 |
| | | D.long019819.01 | 27 | 42 | 9 | 1 | 8 |
| | | D.long019820.01 | 21 | 0 | 21 | 11 | 10 |
| | | D.long019821.01 | 11 | 46 | 10 | 4 | 6 |
| | | D.long019822.01 | 5 | 5 | 4 | 2 | 2 |
| | | D.long019823.01 | 13 | 33 | 19 | 9 | 10 |
| | | D.long019824.01 | 10 | 98 | 16 | 9 | 7 |
| | | D.long019825.01 | 49 | 10 | 90 | 72 | 18 |
| | | D.long019826.01 | 9 | 50 | 62 | 18 | 44 |
| | | D.long019827.01 | 11 | 12 | 4 | 3 | 1 |
| | | D.long019828.01 | 10 | 0 | 16 | 10 | 6 |
| | chr9 | D.long022501.01 | 1 | 0 | 62 | 27 | 35 |
| | | D.long022502.01 | 40 | 178 | 81 | 13 | 68 |
| | | D.long022503.01 | 13 | 7 | 21 | 7 | 14 |
| | | D.long022504.01 | 36 | 150 | 139 | 41 | 98 |
| | | D.long022505.01 | 26 | 12 | 89 | 29 | 60 |
| | | D.long022506.01 | 2 | 5 | 79 | 33 | 46 |
| | | D.long022507.01 | 38 | 18 | 42 | 11 | 31 |
| | | D.long022508.01 | 1 | 7 | 85 | 36 | 49 |
| | | D.long022509.01 | 8 | 19 | 69 | 26 | 43 |
| | | D.long022510.01 | 3 | 0 | 67 | 28 | 39 |
| | | D.long022511.01 | 22 | 13 | 36 | 11 | 25 |
| | | D.long022512.01 | 52 | 0 | 213 | 70 | 143 |
| | | D.long022513.01 | 33 | 63 | 38 | 12 | 26 |
| | | D.long022514.01 | 32 | 118 | 78 | 43 | 35 |
| | | D.long022515.01 | 30 | 0 | 6 | 6 | 0 |
| | | D.long022516.01 | 2 | 29 | 28 | 9 | 19 |
| | | D.long022517.01 | 28 | 60 | 28 | 8 | 20 |
| | | D.long022518.01 | 16 | 48 | 68 | 21 | 47 |
| | | D.long022519.01 | 1 | 2 | 2 | 2 | 0 |
| | | D.long022520.01 | 13 | 11 | 24 | 8 | 16 |
| | | D.long022521.01 | 19 | 28 | 1 | 0 | 1 |

Table S11. SNPs at three genes related to TSS and seed weight .

**Gene ID: D. long019821**

| ID | gene element | Reference | Alternate | BYZ-FJ | CK-FJ | HBP-FJ | FY-FJ | QZB-FJ | JYW-FJ | GSEH-GD | HLGY-GD | YTB-FJ | LQB-FJ | LY-GD | JL-FJ | WLL-FJ | SSCR-GD | JY-GD | SX-GD | HH-GD | HDGY-GD |
|---|---|---|---|---|---|---|---|---|---|---|---|---|---|---|---|---|---|---|---|---|---|
| chr3:18379687 | CDS | A | T | 0 0 | 0 0 | 0 0 | 0 0 | 0 0 | 0 0 | 0 0 | 0 0 | 0 0 | 0 0 | 0 0 | 0 0 | 0 0 | 0 1 | 0 1 | 0 1 | 0 1 | 0 1 |
| chr3:18379727 | CDS | A | G | 0 0 | 0 0 | 0 0 | 0 0 | 0 0 | 0 0 | 0 0 | 0 0 | 0 0 | 0 0 | 0 0 | 0 0 | 0 0 | 0 1 | 0 1 | 0 1 | 0 1 | 0 1 |
| chr3:18380152 | Splice | G | C | 0 0 | 0 0 | 0 0 | 0 0 | 0 0 | 0 0 | 0 0 | 0 0 | 0 0 | 0 0 | 0 0 | 0 0 | 0 0 | 0 1 | 0 1 | 0 1 | 0 1 | 0 1 |
| chr3:18380851 | CDS | A | G | 0 0 | 0 0 | 0 0 | 0 0 | 0 0 | 0 0 | 0 0 | 0 0 | 0 0 | 0 0 | 0 0 | 0 0 | 0 0 | 0 1 | 0 1 | 0 1 | 0 1 | 0 1 |
| chr3:18380866 | CDS | A | G | 0 0 | 0 0 | 0 0 | 0 0 | 0 0 | 0 0 | 0 0 | 0 0 | 0 0 | 0 0 | 0 0 | 0 0 | 0 0 | 0 1 | 0 1 | 0 1 | 0 1 | 0 1 |
| chr3:18380889 | Intron | G | A | 0 0 | 0 0 | 0 0 | 0 0 | 0 0 | 0 0 | 0 0 | 0 0 | 0 0 | 0 0 | 0 0 | 0 0 | 0 0 | 0 1 | 0 1 | 0 1 | 0 1 | 0 1 |
| chr3:18380915 | Intron | A | T | 0 0 | 0 0 | 0 0 | 0 0 | 0 0 | 0 0 | 0 0 | 0 0 | 0 0 | 0 0 | 0 0 | 0 0 | 0 0 | 0 1 | 0 1 | 0 1 | 0 1 | 0 1 |
| chr3:18381202 | Intron | C | A | 0 0 | 0 0 | 0 0 | 0 0 | 0 0 | 0 0 | 0 0 | 0 0 | 0 0 | 0 0 | 0 0 | 0 0 | 0 0 | 0 1 | 0 1 | 0 1 | 0 1 | 0 1 |
| chr3:18381237 | Intron | C | A | 0 0 | 0 0 | 0 0 | 0 0 | 0 0 | 0 0 | 0 0 | 0 0 | 0 0 | 0 0 | 0 0 | 0 0 | 0 0 | 0 1 | 0 1 | 0 1 | 0 1 | 0 1 |
| chr3:18381266 | Intron | T | A | 0 0 | 0 0 | 0 0 | 0 0 | 0 0 | 0 0 | 0 0 | 0 0 | 0 0 | 0 0 | 0 0 | 0 0 | 0 0 | 0 1 | 0 1 | 0 1 | 0 1 | 0 1 |
| chr3:18381657 | Intron | G | T | 0 0 | 0 0 | 0 0 | 0 0 | 0 0 | 0 0 | 0 0 | 0 0 | 0 0 | 0 0 | 0 0 | 0 0 | 0 0 | 0 1 | 0 1 | 0 1 | 0 1 | 0 1 |
| chr3:18381872 | Intron | T | C | 0 0 | 0 0 | 0 0 | 0 0 | 0 0 | 0 0 | 0 0 | 0 0 | 0 0 | 0 0 | 0 0 | 0 0 | 0 0 | 0 1 | 0 1 | 0 1 | 0 1 | 0 1 |
| chr3:18382077 | Intron | A | C | 0 0 | 0 0 | 0 0 | 0 0 | 0 0 | 0 0 | 0 0 | 0 0 | 0 0 | 0 0 | 0 0 | 0 0 | 0 0 | 0 1 | 0 1 | 0 1 | 0 1 | **1 1** |
| chr3:18382363 | Intron | T | A | 0 0 | 0 0 | 0 0 | 0 0 | 0 0 | 0 0 | 0 0 | 0 0 | 0 0 | 0 0 | 0 0 | 0 0 | 0 0 | 0 1 | 0 1 | 0 1 | 0 1 | **1 1** |
| chr3:18382917 | Intron | G | C | 0 0 | 0 0 | 0 0 | 0 0 | 0 0 | 0 0 | 0 0 | 0 0 | 0 0 | 0 0 | 0 0 | 0 0 | 0 0 | 0 1 | 0 1 | 0 1 | 0 1 | **1 1** |
| chr3:18383098 | Intron | T | A | 0 0 | 0 0 | 0 0 | 0 0 | 0 0 | 0 0 | 0 0 | 0 0 | 0 0 | 0 0 | 0 0 | 0 0 | 0 0 | 0 1 | 0 1 | 0 1 | 0 1 | 0 1 |
| chr3:18383184 | Intron | T | C | 0 0 | 0 0 | 0 0 | 0 0 | 0 0 | 0 0 | 0 0 | 0 0 | 0 0 | 0 0 | 0 0 | 0 0 | 0 0 | 0 1 | 0 1 | 0 1 | 0 1 | 0 1 |
| chr3:18383483 | Intron | C | T | 0 0 | 0 0 | 0 0 | 0 0 | 0 0 | 0 0 | 0 0 | 0 0 | 0 0 | 0 0 | 0 0 | 0 0 | 0 0 | 0 1 | 0 1 | 0 1 | 0 1 | 0 1 |
| chr3:18383702 | Intron | G | A | 0 0 | 0 0 | 0 0 | 0 0 | 0 0 | 0 0 | 0 0 | 0 0 | 0 0 | 0 0 | 0 0 | 0 0 | 0 0 | 0 1 | 0 1 | 0 1 | 0 1 | 0 1 |
| chr3:18383849 | Intron | C | T | 0 0 | 0 0 | 0 0 | 0 0 | 0 0 | 0 0 | 0 0 | 0 0 | 0 0 | 0 0 | 0 0 | 0 0 | 0 0 | 0 1 | 0 1 | 0 1 | 0 1 | 0 1 |
| chr3:18383970 | Intron | A | C | 0 0 | 0 0 | 0 0 | 0 0 | 0 0 | 0 0 | 0 0 | 0 0 | 0 0 | 0 0 | 0 0 | 0 0 | 0 0 | 0 1 | 0 1 | 0 1 | 0 1 | 0 1 |
| chr3:18384029 | Intron | A | G | 0 0 | 0 0 | 0 0 | 0 0 | 0 0 | 0 0 | 0 0 | 0 0 | 0 0 | 0 0 | 0 0 | 0 0 | 0 0 | 0 1 | 0 1 | 0 1 | 0 1 | 0 1 |
| chr3:18384183 | Intron | G | C | 0 0 | 0 0 | 0 0 | 0 0 | 0 0 | 0 0 | 0 0 | 0 0 | 0 0 | 0 0 | 0 0 | 0 0 | 0 0 | 0 1 | 0 1 | 0 1 | 0 1 | **1 1** |
| chr3:18384713 | Intron | G | A | 0 0 | 0 0 | 0 0 | 0 0 | 0 0 | 0 0 | 0 0 | 0 0 | 0 0 | 0 0 | 0 0 | 0 0 | 0 0 | 0 1 | 0 1 | 0 1 | 0 0 | 0 1 |
| chr3:18384919 | Intron | G | A | 0 0 | 0 0 | 0 0 | 0 0 | 0 0 | 0 0 | 0 0 | 0 0 | 0 0 | 0 0 | 0 0 | 0 0 | 0 0 | 0 1 | 0 1 | 0 1 | 0 1 | 0 1 |
| chr3:18385255 | Intron | T | C | 0 0 | 0 0 | 0 0 | 0 0 | 0 0 | 0 0 | 0 0 | 0 0 | 0 0 | 0 0 | 0 0 | 0 0 | 0 0 | 0 1 | 0 1 | 0 1 | 0 1 | **1 1** |
| chr3:18385334 | Intron | G | A | 0 0 | 0 0 | 0 0 | 0 0 | 0 0 | 0 0 | 0 0 | 0 0 | 0 0 | 0 0 | 0 0 | 0 0 | 0 0 | 0 1 | 0 1 | 0 1 | 0 1 | 0 1 |
| chr3:18385628 | Intron | A | G | 0 0 | 0 0 | 0 0 | 0 0 | 0 0 | 0 0 | 0 0 | 0 0 | 0 0 | 0 0 | 0 0 | 0 0 | 0 0 | 0 1 | 0 1 | 0 1 | 0 1 | 0 1 |
| chr3:18385671 | Intron | C | T | 0 0 | 0 0 | 0 0 | 0 0 | 0 0 | 0 0 | 0 0 | 0 0 | 0 0 | 0 0 | 0 0 | 0 0 | 0 0 | 0 1 | 0 1 | 0 1 | 0 1 | **1 1** |
| chr3:18385705 | Intron | G | T | 0 0 | 0 0 | 0 0 | 0 0 | 0 0 | 0 0 | 0 0 | 0 0 | 0 0 | 0 0 | 0 0 | 0 0 | 0 0 | 0 1 | 0 1 | 0 1 | 0 1 | 0 1 |
| chr3:18385708 | Intron | C | A | 0 0 | 0 0 | 0 0 | 0 0 | 0 0 | 0 0 | 0 0 | 0 0 | 0 0 | 0 0 | 0 0 | 0 0 | 0 0 | 0 1 | 0 1 | 0 1 | 0 1 | **1 1** |
| chr3:18385893 | Intron | C | G | 0 0 | 0 0 | 0 0 | 0 0 | 0 0 | 0 0 | 0 0 | 0 0 | 0 0 | 0 0 | 0 0 | 0 0 | 0 0 | 0 1 | 0 1 | 0 1 | 0 1 | **1 1** |
| chr3:18386236 | Intron | T | C | 0 0 | 0 0 | 0 0 | 0 0 | 0 0 | 0 0 | 0 0 | 0 0 | 0 0 | 0 0 | 0 0 | 0 0 | 0 0 | 0 1 | 0 1 | 0 1 | 0 1 | **1 1** |
| chr3:18386269 | Intron | G | A | 0 0 | 0 0 | 0 0 | 0 0 | 0 0 | 0 0 | 0 0 | 0 0 | 0 0 | 0 0 | 0 0 | 0 0 | 0 0 | 0 1 | 0 1 | 0 1 | 0 1 | 0 1 |
| chr3:18386527 | Intron | C | T | 0 0 | 0 0 | 0 0 | 0 0 | 0 0 | 0 0 | 0 0 | 0 0 | 0 0 | 0 0 | 0 0 | 0 0 | 0 0 | 0 1 | 0 1 | 0 1 | 0 1 | 0 1 |
| chr3:18386555 | Intron | A | G | 0 0 | 0 0 | 0 0 | 0 0 | 0 0 | 0 0 | 0 0 | 0 0 | 0 0 | 0 0 | 0 0 | 0 0 | 0 0 | 0 1 | 0 1 | 0 1 | 0 1 | **1 1** |
| chr3:18386896 | Intron | G | T | 0 0 | 0 0 | 0 0 | 0 0 | 0 0 | 0 0 | 0 0 | 0 0 | 0 0 | 0 0 | 0 0 | 0 0 | 0 0 | 0 1 | 0 1 | 0 1 | 0 1 | 0 1 |
| chr3:18386914 | Intron | T | G | 0 0 | 0 0 | 0 0 | 0 0 | 0 0 | 0 0 | 0 0 | 0 0 | 0 0 | 0 0 | 0 0 | 0 0 | 0 0 | 0 1 | 0 1 | 0 1 | 0 1 | 0 1 |
| chr3:18387476 | Intron | A | C | 0 0 | 0 0 | 0 0 | 0 0 | 0 0 | 0 0 | 0 0 | 0 0 | 0 0 | 0 0 | 0 0 | 0 0 | 0 0 | 0 1 | 0 1 | 0 1 | 0 1 | **1 1** |
| chr3:18387539 | Intron | A | G | 0 0 | 0 0 | 0 0 | 0 0 | 0 0 | 0 0 | 0 0 | 0 0 | 0 0 | 0 0 | 0 0 | 0 0 | 0 0 | 0 1 | 0 1 | 0 1 | 0 1 | 0 1 |
| chr3:18387641 | CDS | C | T | 0 0 | 0 0 | 0 0 | 0 0 | 0 0 | 0 0 | 0 0 | 0 0 | 0 0 | 0 0 | 0 0 | 0 0 | 0 0 | 0 1 | 0 1 | 0 1 | 0 1 | **1 1** |
| chr3:18387755 | CDS | T | C | 0 0 | 0 0 | 0 0 | 0 0 | 0 0 | 0 0 | 0 0 | 0 0 | 0 0 | 0 0 | 0 0 | 0 0 | 0 0 | 0 1 | 0 1 | 0 1 | 0 1 | 0 1 |
| chr3:18388087 | CDS | G | A | 0 0 | 0 0 | 0 0 | 0 0 | 0 0 | 0 0 | 0 0 | 0 0 | 0 0 | 0 0 | 0 0 | 0 0 | 0 0 | 0 1 | 0 1 | 0 1 | 0 1 | 0 1 |
| chr3:18388347 | Intron | T | C | 0 0 | 0 0 | 0 0 | 0 0 | 0 0 | 0 0 | 0 0 | 0 0 | 0 0 | 0 0 | 0 0 | 0 0 | 0 0 | 0 1 | 0 1 | 0 1 | 0 1 | **1 1** |
| chr3:18388400 | Intron | G | A | 0 0 | 0 0 | 0 0 | 0 0 | 0 0 | 0 0 | 0 0 | 0 0 | 0 0 | 0 0 | 0 0 | 0 0 | 0 0 | 0 1 | 0 1 | 0 1 | 0 1 | 0 1 |
| chr3:18388487 | Intron | T | A | 0 0 | 0 0 | 0 0 | 0 0 | 0 0 | 0 0 | 0 0 | 0 0 | 0 0 | 0 0 | 0 0 | 0 0 | 0 0 | 0 1 | 0 1 | 0 1 | 0 1 | **1 1** |
| chr3:18388518 | Intron | T | C | 0 0 | 0 0 | 0 0 | 0 0 | 0 0 | 0 0 | 0 0 | 0 0 | 0 0 | 0 0 | 0 0 | 0 0 | 0 0 | 0 1 | 0 1 | 0 1 | 0 1 | **1 1** |
| chr3:18388579 | Intron | A | G | 0 0 | 0 0 | 0 0 | 0 0 | 0 0 | 0 0 | 0 0 | 0 0 | 0 0 | 0 0 | 0 0 | 0 0 | 0 0 | 0 1 | 0 1 | 0 1 | 0 1 | 0 1 |
| chr3:18388617 | Intron | T | C | 0 0 | 0 0 | 0 0 | 0 0 | 0 0 | 0 0 | 0 0 | 0 0 | 0 0 | 0 0 | 0 0 | 0 0 | 0 0 | 0 1 | 0 1 | 0 1 | 0 1 | **1 1** |
| chr3:18388628 | Intron | A | G | 0 0 | 0 0 | 0 0 | 0 0 | 0 0 | 0 0 | 0 0 | 0 0 | 0 0 | 0 0 | 0 0 | 0 0 | 0 0 | 0 1 | 0 1 | 0 1 | 0 1 | **1 1** |
| chr3:18388925 | Intron | C | T | 0 0 | 0 0 | 0 0 | 0 0 | 0 0 | 0 0 | 0 0 | 0 0 | 0 0 | 0 0 | 0 0 | 0 0 | 0 0 | 0 1 | 0 1 | 0 1 | 0 1 | 0 1 |
| chr3:18389085 | Intron | A | G | 0 0 | 0 0 | 0 0 | 0 0 | 0 0 | 0 0 | 0 0 | 0 0 | 0 0 | 0 0 | 0 0 | 0 0 | 0 0 | 0 0 | 0 0 | 0 0 | 0 0 | 0 1 |
| chr3:18389103 | Intron | C | A | 0 0 | 0 0 | 0 0 | 0 0 | 0 0 | 0 0 | 0 0 | 0 0 | 0 0 | 0 0 | 0 0 | 0 0 | 0 0 | 0 1 | 0 1 | 0 1 | 0 0 | 0 1 |
| chr3:18389111 | Intron | G | T | 0 0 | 0 0 | 0 0 | 0 0 | 0 0 | 0 0 | 0 0 | 0 0 | 0 0 | 0 0 | 0 0 | 0 0 | 0 0 | 0 1 | 0 1 | 0 1 | 0 1 | 0 1 |
| chr3:18389180 | CDS | A | G | 0 0 | 0 0 | 0 0 | 0 0 | 0 0 | 0 0 | 0 0 | 0 0 | 0 0 | 0 0 | 0 0 | 0 0 | 0 0 | 0 0 | 0 0 | 0 0 | 0 1 | **1 1** |
| chr3:18389234 | CDS | C | G | 0 0 | 0 0 | 0 0 | 0 0 | 0 0 | 0 0 | 0 0 | 0 0 | 0 0 | 0 0 | 0 0 | 0 0 | 0 0 | 0 1 | 0 1 | 0 1 | 0 1 | **1 1** |
| chr3:18389338 | CDS | T | C | 0 0 | 0 0 | 0 0 | 0 0 | 0 0 | 0 0 | 0 0 | 0 0 | 0 0 | 0 0 | 0 0 | 0 0 | 0 0 | 0 1 | 0 1 | 0 1 | 0 1 | **1 1** |
| chr3:18389374 | Promoter | C | G | 0 0 | 0 0 | 0 0 | 0 0 | 0 0 | 0 0 | 0 0 | 0 0 | 0 0 | 0 0 | 0 0 | 0 0 | 0 0 | 0 1 | 0 1 | 0 1 | 0 1 | **1 1** |
| chr3:18389426 | Promoter | T | C | 0 0 | 0 0 | 0 0 | 0 0 | 0 0 | 0 0 | 0 0 | 0 0 | 0 0 | 0 0 | 0 0 | 0 0 | 0 0 | 0 1 | 0 1 | 0 1 | 0 1 | 0 1 |
| chr3:18389437 | Promoter | A | G | 0 0 | 0 0 | 0 0 | 0 0 | 0 0 | 0 0 | 0 0 | 0 0 | 0 0 | 0 0 | 0 0 | 0 0 | 0 0 | 0 1 | 0 1 | 0 1 | 0 1 | 0 1 |
| chr3:18389461 | Promoter | C | G | 0 0 | 0 0 | 0 0 | 0 0 | 0 0 | 0 0 | 0 0 | 0 0 | 0 0 | 0 0 | 0 0 | 0 0 | 0 0 | 0 1 | 0 1 | 0 1 | 0 1 | 0 1 |
| chr3:18389498 | Promoter | T | C | 0 0 | 0 0 | 0 0 | 0 0 | 0 0 | 0 0 | 0 0 | 0 0 | 0 0 | 0 0 | 0 0 | 0 0 | 0 0 | 0 1 | 0 1 | 0 1 | 0 1 | 0 1 |
| chr3:18389556 | Promoter | C | T | 0 0 | 0 0 | 0 0 | 0 0 | 0 0 | 0 0 | 0 0 | 0 0 | 0 0 | 0 0 | 0 0 | 0 0 | 0 0 | 0 1 | 0 1 | 0 1 | 0 1 | 0 1 |
| chr3:18389593 | Promoter | T | C | 0 0 | 0 0 | 0 0 | 0 0 | 0 0 | 0 0 | 0 0 | 0 0 | 0 0 | 0 0 | 0 0 | 0 0 | 0 0 | 0 1 | 0 1 | 0 1 | 0 1 | 0 1 |
| chr3:18389601 | Promoter | G | A | 0 0 | 0 0 | 0 0 | 0 0 | 0 0 | 0 0 | 0 0 | 0 0 | 0 0 | 0 0 | 0 0 | 0 0 | 0 0 | 0 1 | 0 1 | 0 1 | 0 1 | 0 1 |
| chr3:18389722 | Promoter | G | A | 0 0 | 0 0 | 0 0 | 0 0 | 0 0 | 0 0 | 0 0 | 0 0 | 0 0 | 0 0 | 0 0 | 0 0 | 0 0 | 0 1 | 0 1 | 0 1 | 0 1 | 0 1 |
| chr3:18389802 | Promoter | G | T | 0 0 | 0 0 | 0 0 | 0 0 | 0 0 | 0 0 | 0 0 | 0 0 | 0 0 | 0 0 | 0 0 | 0 0 | 0 0 | 0 1 | 0 1 | 0 1 | 0 1 | **1 1** |
| chr3:18389834 | Promoter | T | G | 0 0 | 0 0 | 0 0 | 0 0 | 0 0 | 0 0 | 0 0 | 0 0 | 0 0 | 0 0 | 0 0 | 0 0 | 0 0 | 0 1 | 0 1 | 0 1 | 0 1 | 0 1 |

**Gene ID: D. long019822**

| ID | gene element | Reference | Alternate | BYZ-FJ | CK-FJ | HBP-FJ | FY-FJ | QZB-FJ | JYW-FJ | GSEH-GD | HLGY-GD | YTB-FJ | LQB-FJ | LY-GD | JL-FJ | WLL-FJ | SSCR-GD | JY-GD | SX-GD | HH-GD | HDGY-GD |
|---|---|---|---|---|---|---|---|---|---|---|---|---|---|---|---|---|---|---|---|---|---|
| chr3:18391540 | CDS | G | A | 0_0 | 0_0 | 0_0 | 0_0 | 0_0 | 0_0 | 0_0 | 0_0 | 0_0 | 0_0 | 0_0 | 0_0 | 0_0 | 0_1 | 0_1 | 0_1 | 0_1 | 0_1 |
| chr3:18391690 | CDS | G | A | 0_0 | 0_0 | 0_0 | 0_0 | 0_0 | 0_0 | 0_0 | 0_0 | 0_0 | 0_0 | 0_0 | 0_0 | 0_0 | 0_1 | 0_1 | 0_1 | 0_1 | 0_1 |
| chr3:18392224 | Intron | T | C | 0_0 | 0_0 | 0_0 | 0_0 | 0_0 | 0_0 | 0_0 | 0_0 | 0_0 | 0_0 | 0_0 | 0_0 | 0_0 | 0_1 | 0_1 | 0_1 | 0_1 | **1_1** |
| chr3:18392235 | Intron | T | C | 0_0 | 0_0 | 0_0 | 0_0 | 0_0 | 0_0 | 0_0 | 0_0 | 0_0 | 0_0 | 0_0 | 0_0 | 0_0 | 0_1 | 0_1 | 0_1 | 0_1 | 0_1 |
| chr3:18392382 | Intron | T | C | 0_0 | 0_0 | 0_0 | 0_0 | 0_0 | 0_0 | 0_0 | 0_0 | 0_0 | 0_0 | 0_0 | 0_0 | 0_0 | 0_1 | 0_1 | 0_1 | 0_1 | **1_1** |
| chr3:18392908 | CDS | A | G | 0_0 | 0_0 | 0_0 | 0_0 | 0_0 | 0_0 | 0_0 | 0_0 | 0_0 | 0_0 | 0_0 | 0_0 | 0_0 | 0_0 | 0_0 | 0_0 | 0_0 | 0_1 |
| chr3:18392909 | CDS | T | C | 0_0 | 0_0 | 0_0 | 0_0 | 0_0 | 0_0 | 0_0 | 0_0 | 0_0 | 0_0 | 0_0 | 0_0 | 0_0 | 0_0 | 0_0 | 0_0 | 0_0 | 0_1 |
| chr3:18393109 | Intron | T | G | 0_0 | 0_0 | 0_0 | 0_0 | 0_0 | 0_0 | 0_0 | 0_0 | 0_0 | 0_0 | 0_0 | 0_0 | 0_0 | 0_0 | 0_0 | 0_0 | 0_0 | 0_1 |
| chr3:18393110 | Intron | C | T | 0_0 | 0_0 | 0_0 | 0_0 | 0_0 | 0_0 | 0_0 | 0_0 | 0_0 | 0_0 | 0_0 | 0_0 | 0_0 | 0_0 | 0_0 | 0_0 | 0_0 | 0_1 |
| chr3:18393922 | Promoter | C | G | 0_0 | 0_0 | 0_0 | 0_0 | 0_0 | 0_0 | 0_0 | 0_0 | 0_0 | 0_0 | 0_0 | 0_0 | 0_0 | 0_0 | 0_0 | 0_0 | 0_0 | 0_1 |
| chr3:18393939 | Promoter | T | A | 0_0 | 0_0 | 0_0 | 0_0 | 0_0 | 0_0 | 0_0 | 0_0 | 0_0 | 0_0 | 0_0 | 0_0 | 0_0 | 0_0 | 0_0 | 0_0 | 0_0 | 0_1 |
| chr3:18393961 | Promoter | A | G | 0_0 | 0_0 | 0_0 | 0_0 | 0_0 | 0_0 | 0_0 | 0_0 | 0_0 | 0_0 | 0_0 | 0_0 | 0_0 | 0_1 | 0_1 | 0_1 | 0_0 | **1_1** |
| chr3:18394023 | Promoter | A | T | 0_0 | 0_0 | 0_0 | 0_0 | 0_0 | 0_0 | 0_0 | 0_0 | 0_0 | 0_0 | 0_0 | 0_0 | 0_0 | 0_1 | 0_1 | 0_1 | 0_0 | **1_1** |
| chr3:18394051 | Promoter | G | A | 0_0 | 0_0 | 0_0 | 0_0 | 0_0 | 0_0 | 0_0 | 0_0 | 0_0 | 0_0 | 0_0 | 0_0 | 0_0 | 0_1 | 0_1 | 0_1 | 0_0 | **1_1** |

**Gene ID: D. long019823**

| ID | gene element | Reference | Alternate | BYZ-FJ | CK-FJ | HBP-FJ | FY-FJ | QZB-FJ | JYW-FJ | GSEH-GD | HLGY-GD | YTB-FJ | LQB-FJ | LY-GD | JL-FJ | WLL-FJ | SSCR-GD | JY-GD | SX-GD | HH-GD | HDGY-GD |
|---|---|---|---|---|---|---|---|---|---|---|---|---|---|---|---|---|---|---|---|---|---|
| chr3:18405540 | Promoter | C | T | 0 0 | 0 0 | 0 0 | 0 0 | 0 0 | 0 0 | 0 0 | 0 0 | 0 0 | 0 0 | 0 0 | 0 0 | 0 0 | 0 1 | 0 1 | 0 1 | 0 0 | 0 1 |
| chr3:18405577 | Promoter | T | C | 0 0 | 0 0 | 0 0 | 0 0 | 0 0 | 0 0 | 0 0 | 0 0 | 0 0 | 0 0 | 0 0 | 0 0 | 0 0 | 0 0 | 0 0 | 0 0 | 0 0 | 0 0 |
| chr3:18405608 | Promoter | A | G | 0 0 | 0 0 | 0 0 | 0 0 | 0 0 | 0 0 | 0 0 | 0 0 | 0 0 | 0 0 | 0 0 | 0 0 | 0 0 | 0 0 | 0 0 | 0 0 | 0 0 | 0 0 |
| chr3:18405618 | Promoter | T | C | 0 0 | 0 0 | 0 0 | 0 0 | 0 0 | 0 0 | 0 0 | 0 0 | 0 0 | 0 0 | 0 0 | 0 0 | 0 0 | 0 0 | 0 0 | 0 0 | 0 0 | 0 0 |
| chr3:18405620 | Promoter | C | A | 0 0 | 0 0 | 0 0 | 0 0 | 0 0 | 0 0 | 0 0 | 0 0 | 0 0 | 0 0 | 0 0 | 0 0 | 0 0 | 0 1 | 0 1 | 0 1 | 0 0 | 0 1 |
| chr3:18405629 | Promoter | T | C | 0 0 | 0 0 | 0 0 | 0 0 | 0 0 | 0 0 | 0 0 | 0 0 | 0 0 | 0 0 | 0 0 | 0 0 | 0 0 | 0 0 | 0 0 | 0 0 | 0 0 | 0 0 |
| chr3:18405638 | Promoter | C | T | 0 0 | 0 0 | 0 0 | 0 0 | 0 0 | 0 0 | 0 0 | 0 0 | 0 0 | 0 0 | 0 0 | 0 0 | 0 0 | 0 1 | 0 1 | 0 1 | 0 0 | **1 1** |
| chr3:18405646 | Promoter | T | C | 0 0 | 0 0 | 0 0 | 0 0 | 0 0 | 0 0 | 0 0 | 0 0 | 0 0 | 0 0 | 0 0 | 0 0 | 0 0 | 0 1 | 0 1 | 0 1 | 0 0 | **1 1** |
| chr3:18405671 | Promoter | C | T | 0 0 | 0 0 | 0 0 | 0 0 | 0 0 | 0 0 | 0 0 | 0 0 | 0 0 | 0 0 | 0 0 | 0 0 | 0 0 | 0 1 | 0 1 | 0 1 | 0 0 | 0 1 |
| chr3:18405754 | Promoter | T | A | 0 0 | 0 0 | 0 0 | 0 0 | 0 0 | 0 0 | 0 0 | 0 0 | 0 0 | 0 0 | 0 0 | 0 0 | 0 0 | 0 0 | 0 0 | 0 0 | 0 0 | 0 0 |
| chr3:18405780 | Promoter | C | T | 0 0 | 0 0 | 0 0 | 0 0 | 0 0 | 0 0 | 0 0 | 0 0 | 0 0 | 0 0 | 0 0 | 0 0 | 0 0 | 0 1 | 0 1 | 0 1 | 0 0 | **1 1** |
| chr3:18405976 | Promoter | C | A | 0 0 | 0 0 | 0 0 | 0 0 | 0 0 | 0 0 | 0 0 | 0 0 | 0 0 | 0 0 | 0 0 | 0 0 | 0 0 | 0 0 | 0 0 | 0 0 | 0 0 | 0 0 |
| chr3:18406084 | CDS | T | G | 0 0 | 0 0 | 0 0 | 0 0 | 0 0 | 0 0 | 0 0 | 0 0 | 0 0 | 0 0 | 0 0 | 0 0 | 0 0 | 0 1 | 0 1 | 0 1 | 0 1 | **1 1** |
| chr3:18406198 | Intron | G | A | 0 0 | 0 0 | 0 0 | 0 0 | 0 0 | 0 0 | 0 0 | 0 0 | 0 0 | 0 0 | 0 0 | 0 0 | 0 0 | 0 0 | 0 0 | 0 0 | 0 0 | 0 0 |
| chr3:18406586 | Intron | T | C | 0 0 | 0 0 | 0 0 | 0 0 | 0 0 | 0 0 | 0 0 | 0 0 | 0 0 | 0 0 | 0 0 | 0 0 | 0 0 | 0 1 | 0 1 | 0 1 | 0 0 | **1 1** |
| chr3:18406600 | Intron | C | T | 0 0 | 0 0 | 0 0 | 0 0 | 0 0 | 0 0 | 0 0 | 0 0 | 0 0 | 0 0 | 0 0 | 0 0 | 0 0 | 0 0 | 0 0 | 0 0 | 0 0 | 0 0 |
| chr3:18406720 | Intron | G | T | 0 0 | 0 0 | 0 0 | 0 0 | 0 1 | 0 0 | 0 1 | 0 0 | 0 0 | 0 0 | 0 0 | 0 0 | 0 0 | 0 1 | 0 1 | **1 1** | 0 1 | **1 1** |
| chr3:18406747 | Intron | G | T | 0 0 | 0 0 | 0 0 | 0 0 | 0 0 | 0 0 | 0 0 | 0 0 | 0 0 | 0 0 | 0 0 | 0 0 | 0 0 | 0 1 | 0 1 | 0 1 | 0 0 | 0 1 |
| chr3:18406806 | Intron | T | C | 0 0 | 0 0 | 0 0 | 0 0 | 0 0 | 0 0 | 0 0 | 0 0 | 0 0 | 0 0 | 0 0 | 0 0 | 0 0 | 0 1 | 0 1 | 0 1 | 0 0 | **1 1** |
| chr3:18406861 | Intron | C | A | 0 0 | 0 0 | 0 0 | 0 0 | 0 0 | 0 0 | 0 0 | 0 0 | 0 0 | 0 0 | 0 0 | 0 0 | 0 0 | 0 1 | 0 1 | 0 1 | 0 0 | **1 1** |
| chr3:18406990 | Intron | C | T | 0 0 | 0 0 | 0 0 | 0 0 | 0 0 | 0 0 | 0 0 | 0 0 | 0 0 | 0 0 | 0 0 | 0 0 | 0 0 | 0 1 | 0 1 | 0 1 | 0 0 | **1 1** |
| chr3:18407080 | Intron | T | C | 0 0 | 0 0 | 0 0 | 0 0 | 0 0 | 0 0 | 0 0 | 0 0 | 0 0 | 0 0 | 0 0 | 0 0 | 0 0 | 0 1 | 0 1 | 0 1 | 0 0 | **1 1** |
| chr3:18407551 | Intron | T | C | 0 0 | 0 0 | 0 0 | 0 0 | 0 0 | 0 0 | 0 0 | 0 0 | 0 0 | 0 0 | 0 0 | 0 0 | 0 0 | 0 1 | 0 1 | 0 1 | 0 0 | **1 1** |
| chr3:18407552 | Intron | A | C | 0 0 | 0 0 | 0 0 | 0 0 | 0 0 | 0 0 | 0 0 | 0 0 | 0 0 | 0 0 | 0 0 | 0 0 | 0 0 | 0 1 | 0 1 | 0 1 | 0 0 | **1 1** |
| chr3:18407608 | CDS | T | G | 0 0 | 0 0 | 0 0 | 0 0 | 0 0 | 0 0 | 0 0 | 0 0 | 0 0 | 0 0 | 0 0 | 0 0 | 0 0 | 0 1 | 0 1 | 0 1 | 0 0 | **1 1** |
| chr3:18408488 | Intron | T | C | 0 0 | 0 0 | 0 0 | 0 0 | 0 0 | 0 0 | 0 0 | 0 0 | 0 0 | 0 0 | 0 0 | 0 0 | 0 0 | 0 1 | 0 1 | 0 1 | 0 0 | 0 1 |
| chr3:18408525 | CDS | A | G | 0 0 | 0 0 | 0 0 | 0 0 | 0 0 | 0 0 | 0 0 | 0 0 | 0 0 | 0 0 | 0 0 | 0 0 | 0 0 | 0 0 | 0 0 | 0 0 | 0 0 | 0 0 |
| chr3:18408642 | Intron | G | T | 0 0 | 0 0 | 0 0 | 0 0 | 0 0 | 0 0 | 0 0 | 0 0 | 0 0 | 0 0 | 0 0 | 0 0 | 0 0 | 0 1 | 0 1 | 0 1 | 0 0 | **1 1** |
| chr3:18408658 | Intron | C | G | 0 0 | 0 0 | 0 0 | 0 0 | 0 0 | 0 0 | 0 0 | 0 0 | 0 0 | 0 0 | 0 0 | 0 0 | 0 0 | 0 1 | 0 1 | 0 1 | 0 0 | 0 1 |
| chr3:18408767 | Intron | G | A | 0 0 | 0 0 | 0 0 | 0 0 | 0 0 | 0 0 | 0 0 | 0 0 | 0 0 | 0 0 | 0 0 | 0 0 | 0 0 | 0 1 | 0 1 | 0 1 | 0 0 | **1 1** |
| chr3:18408916 | Intron | C | A | 0 0 | 0 0 | 0 0 | 0 0 | 0 0 | 0 0 | 0 0 | 0 0 | 0 0 | 0 0 | 0 0 | 0 0 | 0 0 | 0 1 | 0 1 | 0 1 | 0 0 | **1 1** |
| chr3:18409019 | Intron | T | G | 0 0 | 0 0 | 0 0 | 0 0 | 0 0 | 0 0 | 0 0 | 0 0 | 0 0 | 0 0 | 0 0 | 0 0 | 0 0 | 0 1 | 0 1 | 0 1 | 0 0 | **1 1** |
| chr3:18409021 | Intron | G | A | 0 0 | 0 0 | 0 0 | 0 0 | 0 0 | 0 0 | 0 0 | 0 0 | 0 0 | 0 0 | 0 0 | 0 0 | 0 0 | 0 1 | 0 1 | 0 1 | 0 0 | 0 1 |
| chr3:18409088 | Intron | G | T | 0 0 | 0 0 | 0 0 | 0 0 | 0 0 | 0 0 | 0 0 | 0 0 | 0 0 | 0 0 | 0 0 | 0 0 | 0 0 | 0 1 | 0 1 | 0 1 | 0 0 | **1 1** |
| chr3:18409249 | Intron | A | G | 0 0 | 0 0 | 0 0 | 0 0 | 0 0 | 0 0 | 0 0 | 0 0 | 0 0 | 0 0 | 0 0 | 0 0 | 0 0 | 0 1 | 0 1 | 0 1 | 0 0 | 0 1 |
| chr3:18410115 | CDS | G | A | 0 0 | 0 0 | 0 0 | 0 0 | 0 0 | 0 0 | 0 0 | 0 0 | 0 0 | 0 0 | 0 0 | 0 0 | 0 0 | 0 1 | 0 1 | 0 1 | 0 0 | 0 1 |
| chr3:18410315 | CDS | G | T | 0 0 | 0 0 | 0 0 | 0 0 | 0 0 | 0 0 | 0 0 | 0 0 | 0 0 | 0 0 | 0 0 | 0 0 | 0 0 | 0 1 | 0 1 | 0 1 | 0 0 | 0 1 |
| chr3:18410317 | CDS | T | A | 0 0 | 0 0 | 0 0 | 0 0 | 0 0 | 0 0 | 0 0 | 0 0 | 0 0 | 0 0 | 0 0 | 0 0 | 0 0 | 0 1 | 0 1 | 0 1 | 0 0 | **1 1** |
| chr3:18411040 | Intron | G | A | 0 0 | 0 0 | 0 0 | 0 0 | 0 0 | 0 0 | 0 0 | 0 0 | 0 0 | 0 0 | 0 0 | 0 0 | 0 0 | 0 1 | 0 1 | 0 1 | 0 0 | **1 1** |
| chr3:18411139 | CDS | G | A | 0 0 | 0 0 | 0 0 | 0 0 | 0 0 | 0 0 | 0 0 | 0 0 | 0 0 | 0 0 | 0 0 | 0 0 | 0 0 | 0 1 | 0 1 | 0 1 | 0 0 | **1 1** |
| chr3:18411493 | CDS | G | A | 0 0 | 0 0 | 0 0 | 0 0 | 0 0 | 0 0 | 0 0 | 0 0 | 0 0 | 0 0 | 0 0 | 0 0 | 0 0 | 0 1 | 0 1 | 0 1 | 0 0 | **1 1** |
| chr3:18411774 | Intron | C | T | 0 0 | 0 0 | 0 0 | 0 0 | 0 0 | 0 0 | 0 0 | 0 0 | 0 0 | 0 0 | 0 0 | 0 0 | 0 0 | 0 1 | 0 1 | 0 1 | 0 0 | 0 1 |
| chr3:18412015 | Intron | G | T | 0 0 | 0 0 | 0 0 | 0 0 | 0 0 | 0 0 | 0 0 | 0 0 | 0 0 | 0 0 | 0 0 | 0 0 | 0 0 | 0 0 | 0 0 | 0 0 | 0 0 | 0 0 |
| chr3:18412018 | Intron | T | A | 0 0 | 0 0 | 0 0 | 0 0 | 0 0 | 0 0 | 0 0 | 0 0 | 0 0 | 0 0 | 0 0 | 0 0 | 0 0 | 0 1 | 0 1 | 0 1 | 0 0 | **1 1** |
| chr3:18412331 | Intron | G | A | 0 0 | 0 0 | 0 0 | 0 0 | 0 0 | 0 0 | 0 0 | 0 0 | 0 0 | 0 0 | 0 0 | 0 0 | 0 0 | 0 1 | 0 1 | 0 1 | 0 0 | **1 1** |
| chr3:18412435 | Intron | T | C | 0 0 | 0 0 | 0 0 | 0 0 | 0 0 | 0 0 | 0 0 | 0 0 | 0 0 | 0 0 | 0 0 | 0 0 | 0 0 | 0 0 | 0 0 | 0 0 | 0 0 | 0 1 |
| chr3:18412436 | Intron | G | T | 0 0 | 0 0 | 0 0 | 0 0 | 0 0 | 0 0 | 0 0 | 0 0 | 0 0 | 0 0 | 0 0 | 0 0 | 0 0 | 0 0 | 0 0 | 0 0 | 0 0 | 0 0 |
| chr3:18412498 | Intron | A | G | 0 0 | 0 0 | 0 0 | 0 0 | 0 0 | 0 0 | 0 0 | 0 0 | 0 0 | 0 0 | 0 0 | 0 0 | 0 0 | 0 1 | 0 1 | 0 1 | 0 0 | 0 1 |
| chr3:18412547 | Intron | G | A | 0 0 | 0 0 | 0 0 | 0 0 | 0 0 | 0 0 | 0 0 | 0 0 | 0 0 | 0 0 | 0 0 | 0 0 | 0 0 | 0 1 | 0 1 | 0 1 | 0 0 | 0 1 |
| chr3:18412689 | Intron | T | A | 0 0 | 0 0 | 0 0 | 0 0 | 0 0 | 0 0 | 0 0 | 0 0 | 0 0 | 0 0 | 0 0 | 0 0 | 0 0 | 0 1 | 0 1 | 0 1 | 0 0 | 0 1 |
| chr3:18412705 | Intron | T | A | 0 0 | 0 0 | 0 0 | 0 0 | 0 0 | 0 0 | 0 0 | 0 0 | 0 0 | 0 0 | 0 0 | 0 0 | 0 0 | 0 1 | 0 1 | 0 1 | 0 0 | 0 1 |
| chr3:18413151 | Intron | C | T | 0 0 | 0 0 | 0 0 | 0 0 | 0 0 | 0 0 | 0 0 | 0 0 | 0 0 | 0 0 | 0 0 | 0 0 | 0 0 | 0 1 | 0 1 | 0 1 | 0 0 | 0 1 |
| chr3:18413585 | Intron | T | A | 0 0 | 0 0 | 0 0 | 0 0 | 0 0 | 0 0 | 0 0 | 0 0 | 0 0 | 0 0 | 0 0 | 0 0 | 0 0 | 0 1 | 0 1 | 0 1 | 0 0 | **1 1** |
| chr3:18413981 | CDS | T | A | 0 0 | 0 0 | 0 0 | 0 0 | 0 0 | 0 0 | 0 0 | 0 0 | 0 0 | 0 0 | 0 0 | 0 0 | 0 0 | 0 1 | 0 1 | 0 1 | 0 0 | 0 1 |
| chr3:18414357 | CDS | G | A | 0 0 | 0 0 | 0 0 | 0 0 | 0 0 | 0 0 | 0 0 | 0 0 | 0 0 | 0 0 | 0 0 | 0 0 | 0 0 | 0 1 | 0 1 | 0 1 | 0 0 | 0 1 |
| chr3:18414406 | CDS | G | A | 0 0 | 0 0 | 0 0 | 0 0 | 0 0 | 0 0 | 0 0 | 0 0 | 0 0 | 0 0 | 0 0 | 0 0 | 0 0 | 0 1 | 0 1 | 0 1 | 0 0 | **1 1** |
| chr3:18414650 | CDS | C | G | 0 0 | 0 0 | 0 0 | 0 0 | 0 0 | 0 0 | 0 0 | 0 0 | 0 0 | 0 0 | 0 0 | 0 0 | 0 0 | 0 1 | 0 1 | 0 1 | 0 0 | 0 1 |
| chr3:18414980 | CDS | T | A | 0 0 | 0 0 | 0 0 | 0 0 | 0 0 | 0 0 | 0 0 | 0 0 | 0 0 | 0 0 | 0 0 | 0 0 | 0 0 | 0 1 | 0 1 | 0 1 | 0 0 | **1 1** |
| chr3:18415364 | CDS | C | A | 0 0 | 0 0 | 0 0 | 0 0 | 0 0 | 0 0 | 0 0 | 0 0 | 0 0 | 0 0 | 0 0 | 0 0 | 0 0 | 0 1 | 0 1 | 0 1 | 0 0 | 0 1 |
| chr3:18415495 | CDS | G | C | 0 0 | 0 0 | 0 0 | 0 0 | 0 0 | 0 0 | 0 0 | 0 0 | 0 0 | 0 0 | 0 0 | 0 0 | 0 0 | 0 1 | 0 1 | 0 1 | 0 0 | 0 1 |
| chr3:18415619 | CDS | C | T | 0 0 | 0 0 | 0 0 | 0 0 | 0 0 | 0 0 | 0 0 | 0 0 | 0 0 | 0 0 | 0 0 | 0 0 | 0 0 | 0 1 | 0 1 | 0 1 | 0 0 | 0 1 |
| chr3:18415852 | CDS | G | C | 0 0 | 0 0 | 0 0 | 0 0 | 0 0 | 0 0 | 0 0 | 0 0 | 0 0 | 0 0 | 0 0 | 0 0 | 0 0 | 0 1 | 0 1 | 0 1 | 0 0 | 0 1 |
| chr3:18415855 | CDS | G | A | 0 0 | 0 0 | 0 0 | 0 0 | 0 0 | 0 0 | 0 0 | 0 0 | 0 0 | 0 0 | 0 0 | 0 0 | 0 0 | 0 1 | 0 1 | 0 1 | 0 0 | **1 1** |
| chr3:18416276 | CDS | T | C | 0 0 | 0 0 | 0 0 | 0 0 | 0 0 | 0 0 | 0 0 | 0 0 | 0 0 | 0 0 | 0 0 | 0 0 | 0 0 | 0 1 | 0 1 | 0 1 | 0 0 | **1 1** |
